# Genotype and environment shape gene regulatory variation in switchgrass

**DOI:** 10.64898/2026.11.11.765962

**Authors:** Colette S. Berg, Caio Guilherme Pereira, Avinash Sreedasyam, John Lovell, Xiaoyu Weng, Jason Bonnette, Yuko Yoshinaga, Chris Daum, Anna Lipzen, Jeremy Schmutz, Thomas E. Juenger, David B. Lowry

## Abstract

Local adaptation is contingent on genotype-by-environment interactions, which are often mediated by variation in gene expression. The causes and consequences of variation in gene expression are complex, and they are rarely evaluated under the multi-dimensional environmental regimes that occur in the field. Here, we investigate switchgrass gene expression variation in contrasting sites in Michigan and Texas. We examined differentially expressed genes between upland and lowland switchgrass genotypes and found that genes with genotype x environment (GxE) interactions were enriched for GO terms related to phosphorus metabolism. After mapping expression quantitative trait loci (eQTL) in a switchgrass diversity panel in both sites, we compared distinct categories of eQTL to investigate their effects on gene expression in contrasting environments and their evolutionary history. Cis regulatory variants are more likely to have consistent effects across sites, while trans regulatory variants were more likely to have site-specific effects, and are more likely to underlie changes in gene expression between sites. Lower minor allele frequency in trans regulatory variants is consistent with stronger purifying selection. Furthermore, site-specific regulatory variants exhibited intermediate minor allele frequencies, consistent with balancing selection or soft sweeps. Finally, motivated by enrichment for phosphorus-related genes exhibiting differential expression between sites, we conducted a GWAS for leaf phosphorus concentration in the diversity panel. By overlaying the GWAS results with the eQTL map, we characterized cis and trans regulation of candidate genes for leaf P concentration in Michigan and Texas. Together, this work provides novel insights on the molecular basis and evolutionary history of GxE interactions in switchgrass.

## Introduction

As sessile organisms, plants cannot migrate under adverse conditions, and thus rely on local adaptation and plasticity, fine-tuning traits to their surrounding environment (Leimu and Fischer 2008; Anderson et al. 2013; Lascoux et al. 2016). Local adaptation is a type of genotype-by-environment (GxE) interaction in which a genotype expresses distinct phenotypes depending on external conditions, and fitness is higher in the home environment compared to a foreign one (Kawecki and Ebert 2004; Savolainen et al. 2013). Understanding the mechanistic basis of GxE interactions is a fundamental goal of evolutionary genetics. Gene expression variation is an important component of GxE: environmental cues influence gene expression, and the resulting expression variation shapes phenotypes and fitness (López-Maury et al. 2008; Boye et al. 2024). Gene expression variation is an important component of rapid adaptation to new environments (Morris et al. 2014; Ghalambor et al. 2015; Mack et al. 2018; Campbell-Staton et al. 2021) and maintenance of fitness in fluctuating environments (Acar et al. 2008; Gracey et al. 2008; Hamann et al. 2021). By investigating how environment and genotype shape traits, expression and gene regulatory variation, we can better understand the molecular basis and evolutionary history of plant variation in diverse environments.

Gene expression variation among contrasting genotypes and environments can be examined through complementary lenses. Quantifying differential expression (DE) in genes and modules of co-expressed genes in contrasting environments can reveal the molecular pathways and mechanisms enabling plants to cope with biotic and abiotic stresses (Matters and Scandalios 1986; Howarth and Ougham 1993; Reinbothe et al. 1994; Hazen et al. 2003; Chinnusamy et al. 2007; Jamil et al. 2011). Furthermore, mapping the variants regulating gene expression, and examining differences in relationships between genotype and gene expression in contrasting sites can reveal the specific mechanisms underlying GxE interactions. Gene expression is genetically controlled by variants in two categories: cis regulatory variants, which affect the expression of a nearby gene, and trans regulatory variants, which influence the expression of distant genes. Regulatory variants can be mapped using an expression genome wide association study (eGWAS) approach, which involves scanning for relationships between SNPs and gene expression (Gilad et al. 2008) or through allele-specific expression (ASE) analyses in F_1_ crosses between divergent parents (Cowles et al. 2002; Yan et al. 2002). eGWAS is limited by population structure and statistical power, but is effective for mapping cis and trans eQTLs to the single nucleotide polymorphism (SNP) level within populations (Korte and Farlow 2013).

Different classes of regulatory variants likely experience distinct evolutionary pressures. For example, trans eQTL are expected to be under stronger purifying selection than cis eQTL because they pleiotropically affect the expression of several genes at once (McGuigan et al. 2014). This prediction is empirically supported in *Capsella grandiflora* (Josephs et al. 2020) and *Mimulus guttatus* (Brown and Kelly 2022): in both systems, the allele frequency spectrum of trans regulatory variants is skewed towards rare alleles, suggesting a history of purifying selection. However, less is known about the difference in selection history between static eQTL, which have consistent effects across conditions, and dynamic eQTL, whose effects on expression exhibit GxE interactions (Ackermann et al. 2013). In general, cis regulatory variants tend to be more static, while trans regulatory variants are more likely to have context-specific effects (Grundberg et al. 2012; Cubillos et al. 2014; Snoek et al. 2012; Snoek et al. 2017; Lovell et al. 2018; Verta and Jones 2019; Ballinger et al. 2023, but see Ackermann et al. 2013). Recent findings from the immunology literature suggest that eQTL that respond to immune stimulation are under more positive selection compared to static eQTL (Nédélec et al. 2016; Quach et al. 2016; Kim-Hellmuth et al. 2017; Harrison et al. 2019; Lea et al. 2022). However, *Arabidopsis* genes with environment-specific expression tend to be under weaker selection (Roberts and Josephs 2023), and switchgrass genes with plastic expression in response to drought do not exhibit signatures of positive selection (Bhaskara et al. 2023). Prior research has mapped cis and trans regulatory responses to stress in diverse organisms, from mice (Chow et al. 2015) to corals (Bellantuono et al. 2012) to plants (Lowry et al. 2013; Gould et al. 2018; Lovell et al. 2018; Liu et al. 2020; Wei et al. 2024). However, fewer studies have conducted eQTL mapping in contrasting common garden environments, where plants integrate and respond to many environmental factors simultaneously (see Wade et al. 2022). Paired common garden experiments are an ideal context to map both static and site-specific cis and trans eQTL and compare their selective histories.

A promising approach for understanding the consequences of environmentally dependent gene regulation is to examine its relationship with physiological traits that are also sensitive to environmental variation. The plant ionome reflects nutrient acquisition, transport, and homeostasis, all of which are shaped by genetic and environmental factors (Salt et al. 2008; Baxter 2009). Genetic variation in elemental accumulation frequently exhibits GxE interactions (Fikas et al. 2019), making ionomic elemental traits useful for studying the consequences of regulatory variation. One of the most critical plant elements is phosphorus, which is an essential nutrient for plant metabolic functions, being a structural component of nucleic acids and membrane lipids and having a key role in energy transfer (Shen et al. 2011, Marschner et al. 2012, Lambers and Olivera 2019) yet the quantitative genetics underlying variation in leaf phosphorus concentration remains understudied (Guilherme Pereira and Des Marais 2020). Phosphorus (P) concentration in leaves varies along a spectrum, with P-rich leaves generally maintaining higher photosynthetic rates but being more susceptible to herbivory and other types of damage (Wright et al. 2004). Leaf P concentration is heritable and has been mapped to major quantitative trait loci in *Arabidopsis* (Bentsink et al. 2003; Vreugdenhil et al. 2004; Waters and Grusak 2008; Ghandilyan et al. 2009; Prinzenberg et al. 2010), *Brassica rapa* (Zhao et al. 2007), rice (Norton et al. 2010) and switchgrass (Zhang et al. 2021). Within-plant phosphorus dynamics are strongly influenced by the environment: inorganic phosphorus availability is determined by rock weathering, but phosphorus must be converted into its organic form to be used by the plant (Ruttenberg 2003). Environmental factors such as soil pH and texture in tandem with microbial factors within the rhizosphere shape a plant’s ability to access bioavailable phosphorus (Shen et al. 2011). Furthermore, increased P improves plant responses to both abiotic and biotic stress (Khan et al. 2023), acting as a critical fulcrum in the balance between nutrition and immunity. A plant’s phosphorus requirements are dynamic, shifting throughout development, seasonal changes, and microbial factors. Thus, leaf P concentration frequently exhibits GxE interactions (Veley et al. 2017; Fikas et al. 2019; Zhang et al. 2021). Leaf P concentration is an essential, environmentally responsive trait and serves as a powerful lens through which to investigate GxE interactions in plants.

Switchgrass (*Panicum virgatum*) is a perennial grass native to North America that has emerged as a biofuel and bioproducts crop due to its high biomass productivity, low resource requirements, and ability to grow on marginal lands (Parrish and Fike 2005; Casler et al. 2011; Casler 2012; Parrish et al. 2012) and has served as a model system for understanding the evolution of plant ecotypes for > 70 years (McMillan 1959). Switchgrass has high nucleotide diversity (Grabowski et al. 2014) and two divergent ecotypes: upland, which is cold hardy and of short stature, and lowland, which is taller, drought tolerant, and more resistant to disease (McMillan 1959; Zhang et al. 2011; Morris et al. 2016). In addition to these divergent upland and lowland ecotypes, switchgrass exhibits near-continuous, climate-adapted variation for many traits across its range (Lovell et al. 2021). Both GWAS and QTL approaches have revealed candidate loci and genes underlying switchgrass trait variation, many of which exhibit GxE interactions (Grabowski et al. 2014; Lowry et al. 2019; Lovell et al. 2021; Zhang et al. 2022; MacQueen et al. 2024). In particular, leaf P content in switchgrass is heritable, has been mapped to major quantitative trait loci, and exhibits GxE interactions in Michigan and Texas common gardens (Zhang et al. 2021). Switchgrass’ diversity, its broad native range, and excellent genomic resources make it a great system to investigate how both genotype and environment shape gene regulatory variation.

Here, we integrate differential expression, and gene co-expression network analysis in contrasting genotypes and habitats with expression GWAS to investigate the impact of genotype and environment on switchgrass regulatory variation, using expression data from leaves. We address the following questions: (1) Which traits, genes, and gene modules exhibit significant GxE interactions? (2) How does cis and trans regulation of gene expression vary between switchgrass populations and environments? (3) How has selection shaped the evolution of cis and trans eQTL? (4) Which candidate genes are related to variation in leaf P concentration, and how does regulation of expression of candidates vary between sites? This work reveals how regulation of gene expression in leaves mediates GxE interactions, identifies potential gene pathways and mechanisms involved in key switchgrass traits, and tests theoretical predictions about the evolution of cis and trans regulatory variation.

## Results

### GxE genes are involved in phosphorus metabolism

To understand how genotype and environment shape trait and expression variation in switchgrass, we measured leaf traits and collected RNA from clonal replicates of lowland (AP13) and upland (DAC6) switchgrass genotypes reared in common gardens in Michigan and Texas. The common gardens vary substantially in temperature, precipitation, and soil properties (Lovell et al. 2021; Edwards et al. 2023), and the study primarily focused on genes whose expression exhibited significant genotype-by-environment interactions.

Switchgrass gene expression was significantly shaped by genotype, habitat, and their interaction. Of the 32889 genes surveyed, the expression of 12260 (37.2%) was significantly shaped by genotype, 6458 (19.6%) was significantly shaped by environment, and 3165 (9.6%) exhibited significant GxE interaction (Fig. 1a). Of these 3165 GxE genes, 1434 were variance-changing – the change in gene expression between habitats was in the same direction in AP13 and DAC6, with only the degree of change varying (Table S1). The remaining 2125 genes were rank-changing, where AP13 and DAC6 exhibited crossed reaction norms for gene expression between the two sites (Table S1). We also identified 5127 genes whose expression was significantly shaped by genotype and site separately, but whose GxE interaction term was not significant (Table S2).

**Figure 1.**
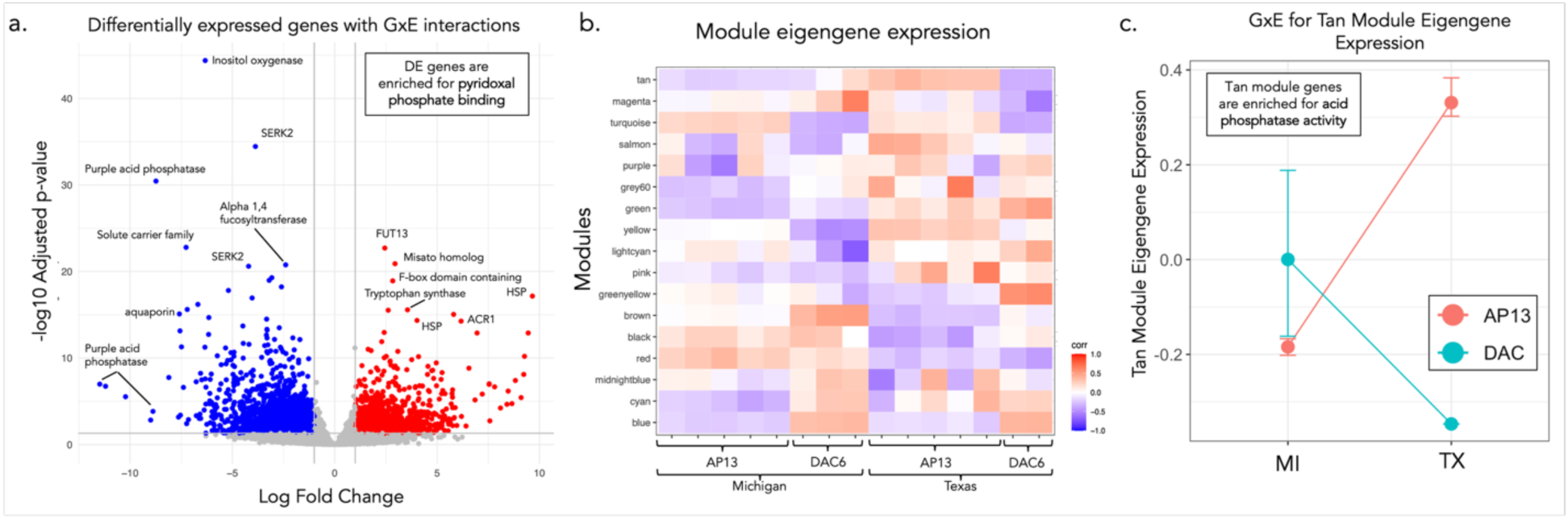
Switchgrass genes with GxE interactions are enriched for phosphorus-related GO terms. a) Volcano plot of genes whose expression is shaped by site, genotype, and their interaction, with select top differentially expressed genes annotated with their function. Pyridoxal phosphate binding is the top GO enrichment term in the GxE genes. b) Module eigengene expression for coexpressed gene modules identified by WGCNA. c) Reaction norm for tan module eigengene expression in AP13 and DAC6 genotypes in Michigan and Texas. Genes within the tan module are significantly enriched for acid phosphatase activity.

Motivated by the enrichment for phosphorus-related terms in the GxE genes, we overlaid the DE genes with previously mapped quantitative trait loci for leaf P in these switchgrass lines (Zhang et al. 2021), and found that many DE genes were within previously mapped leaf P concentration QTL: 2255 (18%) of genotype DE genes, 1638 (25%) of site DE genes, and 465 (15%) of GxE DE genes were located within leaf P QTL intervals (Table S1).

The top GO term in the genes with GxE effects was pyridoxal phosphate binding (Fig 1a, Fig. S1). Additional GO terms enriched in the GxE genes are related to metabolism, transaminase and kinase activity, aromatic amino acid biosynthetic processes, and the pentose-phosphate shunt (Fig. S1). The 3615 significant GxE DE genes included 17 acid phosphatase genes (16 upregulated, 1 downregulated), including 11 purple acid phosphatase genes. Acid phosphatases are involved in phosphorus uptake and metabolism, and purple acid phosphatases specifically mediate responses to oxidative stress (Tran et al. 2010). The DE gene with the most significant GxE interaction was an inositol oxygenase (Fig 1a, Table S1), which regulates the availability of myo-inositol, the molecular precursor to phytic acid, the storage molecule for phosphorus within plant cells (Siddique et al. 2014; Kurita et al. 2017; Poirier et al. 2022). Pyridoxal phosphate binding was the top enriched GO term in the GxE DE genes (Fig. S1). Pyridoxal 5’phosphate is the active form of vitamin B6, an essential metabolite in all organisms that plays a role in plant defense against oxidative stress (Snell and Haskell 1970; Havaux et al. 2009). Acid phosphatases mediate pyridoxal phosphate homeostasis by removing the phosphate group from larger molecules, altering the inorganic phosphate availability within the cell (Huang et al. 2012). The genes exhibiting a significant relationship to site and genotype (but not GxE) were not enriched for phosphorus related GO terms – rather, these genes were enriched for GO terms related to carbohydrate binding, oxidoreductase activity, NAD binding, steroid biosynthetic process, NAD+ activity, metalloendopeptidase activity, and starch binding (Fig. S2). All significant GxE DE genes and their functions can be found in Table S1.

We mapped gene coexpression modules for AP13 and DAC6 in Michigan and Texas, identifying 19 modules. Two of the modules were only up-regulated in one individual and downregulated in all others and were removed from subsequent analysis. Three modules were significantly associated with genotype, seven were significantly associated with site, and three modules were significantly associated with genotype, site, and their interaction (Fig. 1b, Table S3). The tan module had the strongest GxE interaction (p = 1.4e-05, Fig. 1c, Table S3a) and was significantly enriched for two GO terms related to phosphorus metabolism, specifically acid phosphatase activity and phosphoric diester hydrolase activity (Fig. S3). Top hub genes in the tan module included two glycerophosphocoline phosphodiesterases, a purple acid phosphatase gene, and a 4-phytase gene (Table S4), all of which are involved in phosphorus metabolism, specifically hydrolyzing bonds of organic phosphate to make orthophosphate (Pi) available to cells (Wodzinski and Ullah 1996; van der Rest et al. 2002; Vance et al. 2003). GO enrichment plots and top hub genes for all modules can be found in the Supplemental (Fig. S3 – S18, Table S4).

### Site-specific and static eQTL shape cis and trans regulation of switchgrass expression variation across sites

To understand how genetic and environmental variation shape regulation of gene expression for genotypes derived from locations across the species range of switchgrass, we conducted an expression genome wide association study (eGWAS) using RNA sequence and whole genome sequence from the switchgrass diversity panel. The whole genome sequence was first presented in Lovell et al. 2021; the gene expression data was first presented in Izquierido et al. 2026. Because the diversity panel is highly structured by population (Lovell et al. 2021), we conducted separate eGWAS scans for the Midwest, Atlantic, and Gulf populations. For each population, we scanned separately for variants significantly affecting gene expression at the MI site and TX site. Then, we scanned for GxE eQTL by mapping variants significantly associated with the difference in normalized gene expression between MI and TX. Scanning for variants associated with the difference between expression between the two sites allows us to effectively map the GxE component of the reaction norm for gene expression (Aubin-Horth and Renn 2009). To complement the diversity panel cis eQTL mapping, we also identified cis regulated genes through allele specific expression analyses in AP13xDAC6 F_1_’s (details in Supplementary Material).

Because the DE genes between upland and lowland lines in Michigan and Texas were enriched for phosphorus activity (Fig. 1a), we conducted GO term enrichment analysis of the top 10% of diversity panel GxE genes, to assess whether they were enriched for similar functions. We did not find enrichment for phosphorus-related GO terms; rather, the top GxE genes were primarily enriched for terms related to metabolism and RNA processing (Figs. S15 – S17).

eQTL mapping revealed 6968 – 38517 cis eQTL and 18881 – 208240 trans eQTL across populations and sites (Fig. 2a, Fig. S19-S20, Table S5). Cis and trans regulation was common throughout the genome: across populations and sites, 20 – 50 % of all genes were regulated in cis, and 39 – 88% of genes were regulated in trans by at least one variant (Table S5). We also identified trans eQTL hotspots – variants significantly associated with the expression of > 50 genes – in all three subpopulations (Fig. 2a, Fig. S19-S20, Table S6).

**Figure 2.**
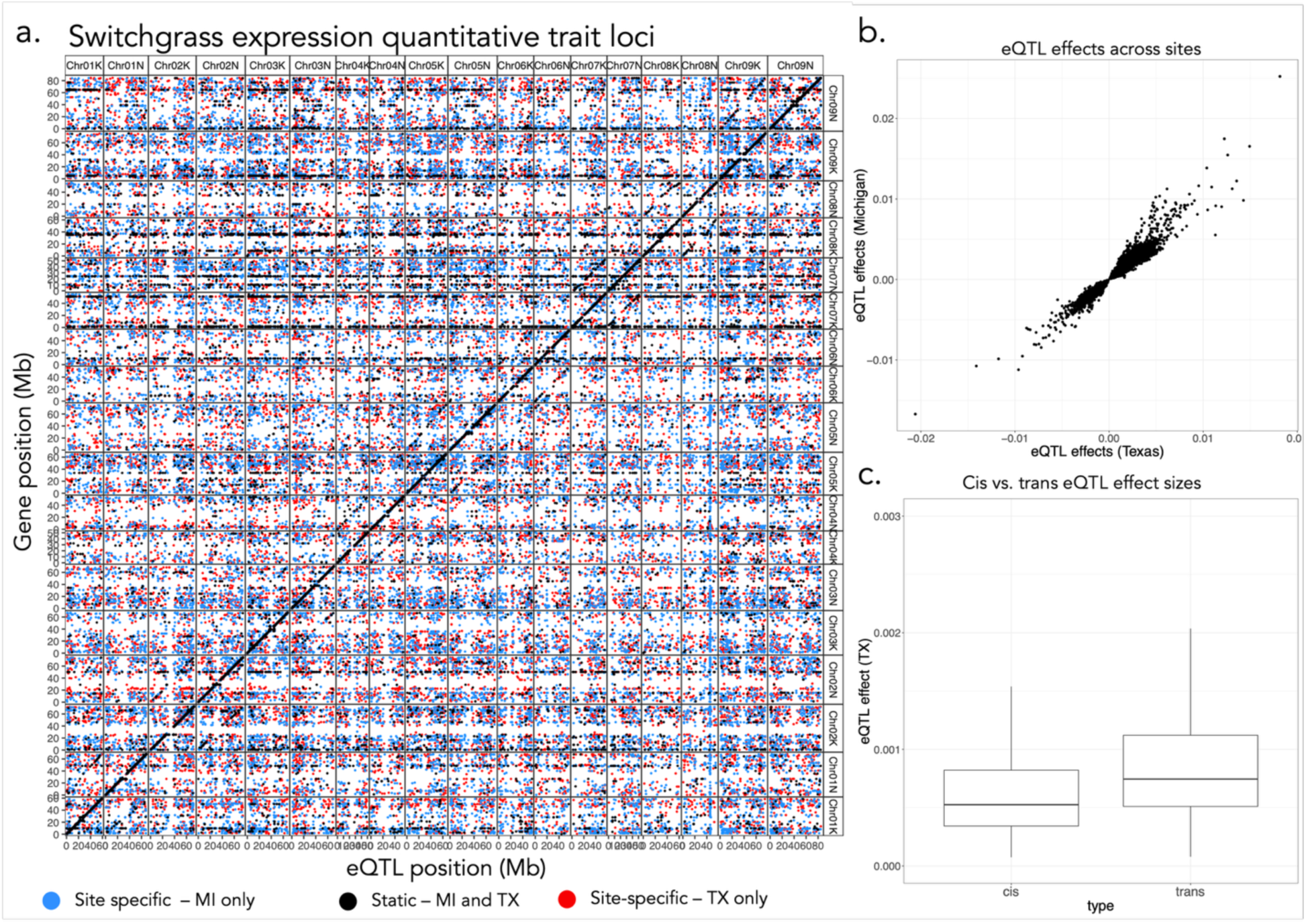
Static and site-specific cis and trans expression quantitative trait loci (eQTL) shape regulation of gene expression in the switchgrass diversity panel. a. Map of eQTL in the Atlantic population of the switchgrass diversity panel. x-axis is the variant position throughout the genome; y-axis is position of the gene regulated. Points along the diagonal represent cis eQTL. b) Relationship between eQTL effect on gene expression at the TX site and eQTL effects on gene expression at the at the MI site. Because of the strong correlation between eQTL effects in TX and MI, we consider eQTL mapped in both sites to have static effects across environments. c) Comparison of the absolute effect size (beta) of cis and trans eQTL on gene expression output by MatrixEQTL. cis and trans eQTL effect sizes were compared with a t-test (trans eQTL beta mean = 0.0006, cis eQTL beta mean = 0.0001, p < 2.2e-16).

We refer to eQTL mapped only in one site (MI or TX) as site-specific eQTL (Fig. 2a, Table S5). We acknowledge that some site-specific QTL may be present in both sites but only meet the significance threshold in one site due to differences in the variant’s effect on gene expression in the two sites. In the eQTL that were mapped in both sites, effect size and direction of eQTL effects in TX and MI were strongly correlated in all three populations (Midwest R^2^ = 0.89, Gulf R^2^ = 0.94, Atlantic R^2^ = 0.95, p < 1e-16 for all populations, Fig. 2b, Fig. S21). We did not find any rank-changing eQTL, i.e. those with opposite direction of effects on gene expression in TX vs. MI (Fig. 2b, Fig, S21). Therefore, we consider the eQTL mapped in both sites to be static eQTL, because they have generally consistent effects on gene expression across sites. Overall, cis eQTL had lower effect size (beta) than trans eQTL across sites and populations (p < 1e-16 for all t-tests between betas for cis and trans eQTL, Fig. 2c, Fig. S22). We did not find any trans eQTL hotspots that were mapped in both sites; rather, all hotspots were site-specific (Table S6).

Switchgrass is a tetraploid with disomic inheritance, and eQTL mapping provided the opportunity to investigate gene regulation within and between sub-genomes. Results and methodology details for the sub-genome comparisons are available in the Supplementary Material (Supplementary Text, Fig. S23, S24)

### cis eQTL tend to have stable effects across space, while trans eQTL are more dynamic

GxE eQTL were those that were significantly associated with a change in gene expression between sites (Fig. 3a, Figs S25-S26). In contrast to the single-site analyses reported above (Fig. 2, Figs S25-S26), in which there were approximately twice as many cis eQTL as trans eQTL, we detected substantially more trans GxE eQTL than cis GxE eQTL (Fig. 3b). This suggests that trans eQTL are more likely to have dynamic effects across space. We also examined cis regulation of genes in contrasting sites by quantifying allele-specific expression (ASE) in AP13xDAC6 F_1_’s and found that 87.7 – 90.3 % of ASE genes had bias in the same direction in both Michigan and Texas. This finding further supports the relative stability of cis regulation across contrasting sites. Results and methodology details for the ASE analyses are available in the Supplementary Material (Supplementary Text, Fig. S27, Table S7).

**Figure 3.**
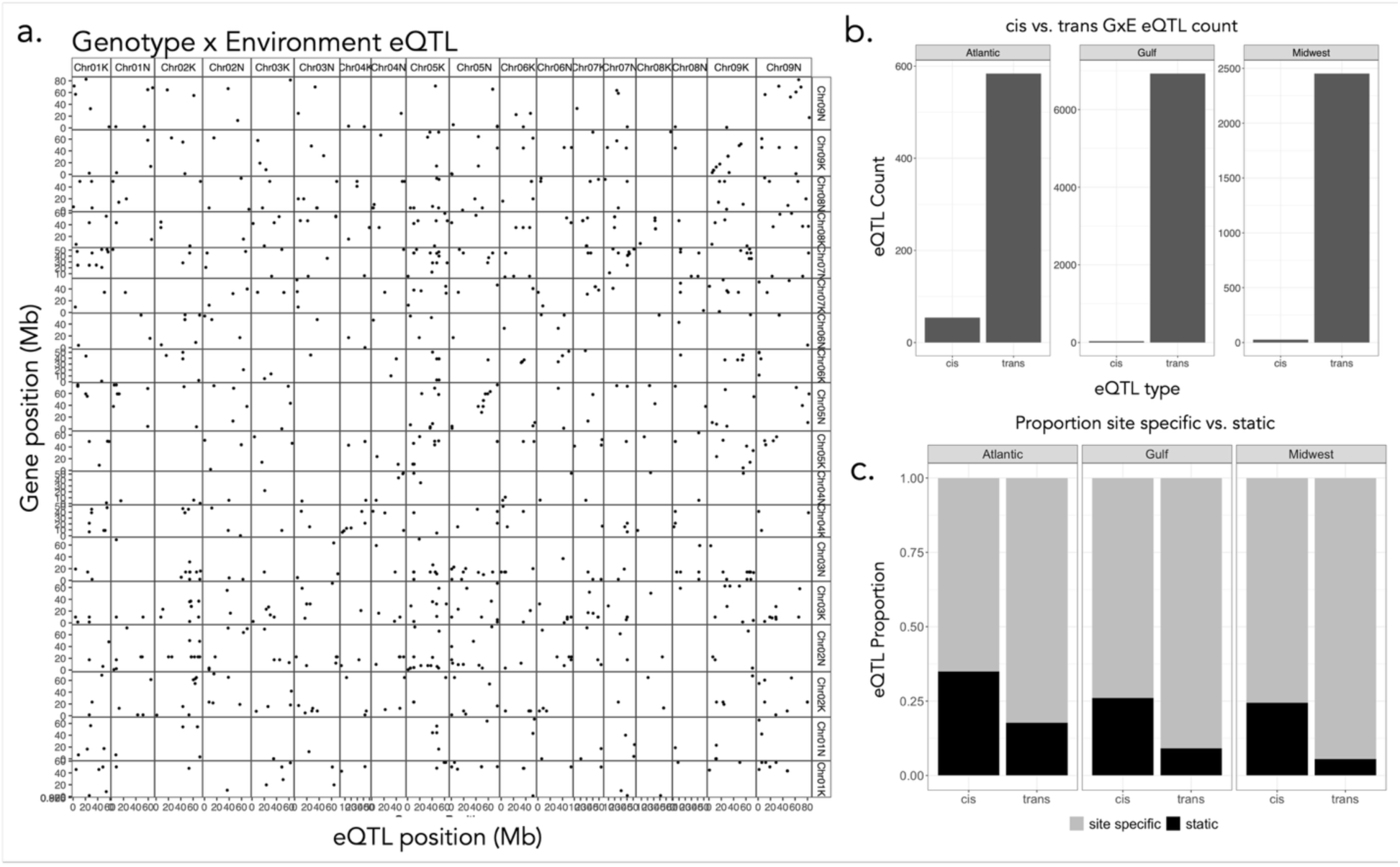
trans eQTL have more dynamic effects across space than cis eQTL. a) Map of expression quantitative trait loci where the variant significantly predicted a change in expression between Michigan and Texas, in the Atlantic population of the diversity panel. x-axis is the variant position throughout the genome; y-axis is position of the gene regulated. Points along the diagonal represent cis eQTL. b) Relative proportion of cite-specific eQTL (those mapped in only MI or only TX) and static eQTL (those mapped in both sites) among cis and trans eQTL in each population of the diversity panel. c) Counts of cis and trans GxE eQTL in each population of the diversity panel.

This tendency for trans eQTL to have more dynamic effects than cis eQTL was reinforced in our comparisons of the site-specific and static eQTL. In each population, the majority of both cis and trans eQTL were site-specific (Fig 2a, Fig. 3c, Table S5). However, a greater proportion of trans eQTL were site-specific compared to cis eQTL (Fig. 3c, Table S5), suggesting that cis regulation of gene was more consistent across sites. Static eQTL generally had greater effect sizes than the site-specific eQTL (Figure S28), and it is possible that the site-specific eQTL were those that do not have strong enough effects to be detected in both environments. However, this difference in effect size does not explain the increased proportion of static cis eQTL compared to trans eQTL, since the cis eQTL generally have lower effect sizes than the trans eQTL (Fig. 2c).

### The site frequency spectrum of trans eQTL is consistent with purifying selection

Because cis and trans eQTL have distinct mechanisms of influencing gene expression, we expect them to have distinct evolutionary histories. To investigate differences in signatures of selection between classes of regulatory variants, we compared the allele frequency spectra of cis and trans eQTL. In all populations and gardens, we found that trans regulatory variants consistently have lower minor allele frequencies (MAF) than cis regulatory variants (p < 1e-16 for all comparisons, Fig. 4a, Fig. S28-S29, Table S8). However, trans eQTL also tend to have larger effect sizes than cis eQTL (Fig. 2c, Fig. S22), and MAF is negatively associated with effect size – i.e., eQTL with lower MAF tend to have higher absolute effect size (p < 2e-16 for all populations and sites, Fig. S30). To investigate whether this difference in allele frequencies between cis and trans eQTL was shaped by differences in effect size, we compared MAFs between cis and trans eQTL in the top quartile of effect sizes and found the same pattern (p < 2.2e-16). Lower minor allele frequencies in the trans eQTL are consistent with stronger purifying selection acting on trans acting regulatory variants (Barton and Keightley 2002).

**Figure 4.**
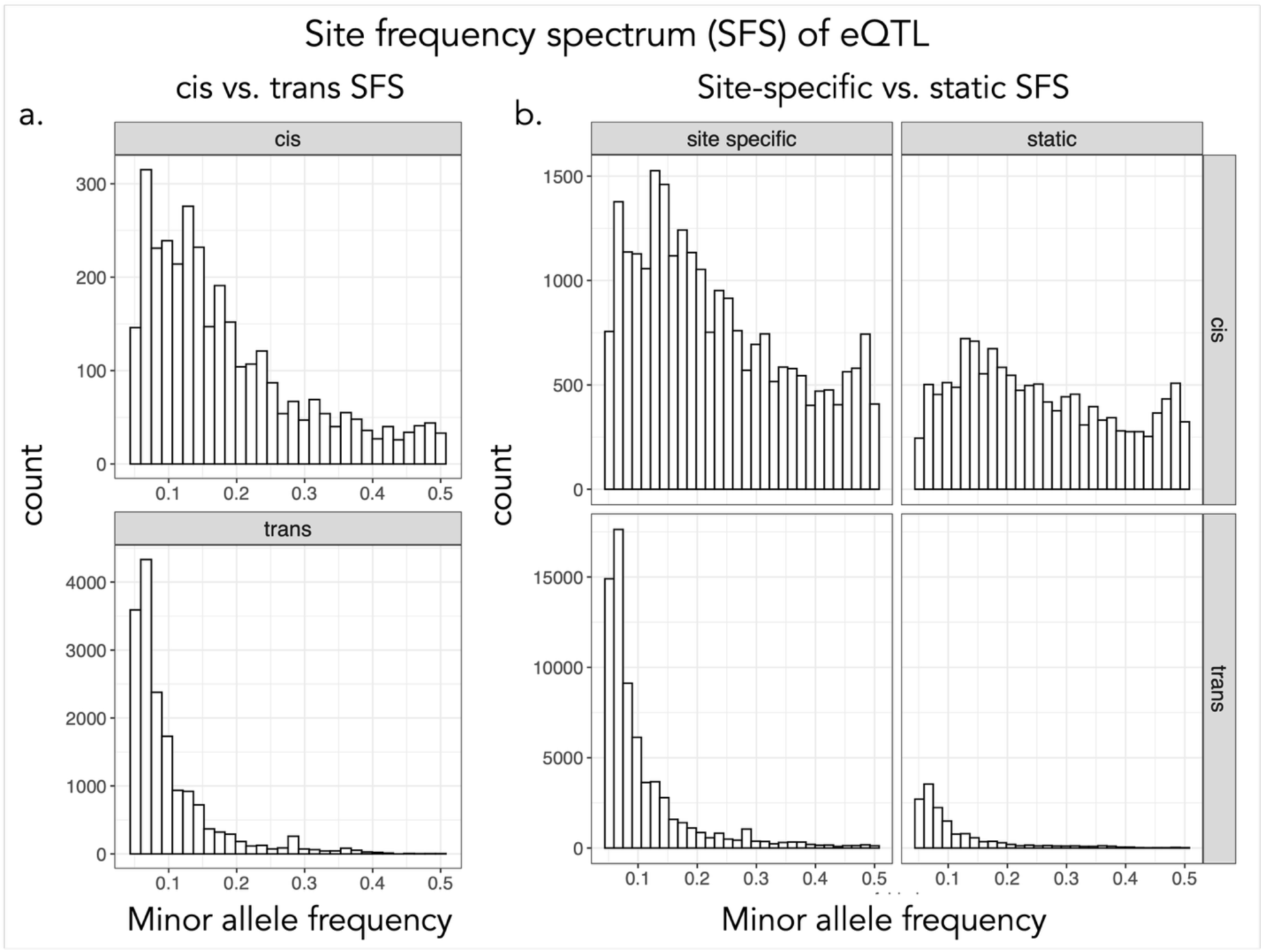
Site frequency spectrum comparisons of eQTL. a) Site frequency spectrum of cis and trans eQTL in the Atlantic population of the diversity panel, with eQTL effects in the top quartile. b) Site frequency spectrum of cis and trans eQTL that are site-specific (mapped in only TX or MI) and static (mapped in both TX and MI) in the Atlantic population of the switchgrass diversity panel.

### Static eQTL have more intermediate minor allele frequencies than site-specific eQTL

We next compared allele frequency spectra of static eQTL – those that were mapped in both MI and TX – and site-specific eQTL, which were mapped only in MI or only in TX. Static eQTL have intermediate allele frequencies compared to site-specific eQTL (p < 2.2e-16 for all comparisons, Fig. 4b, Fig. S28-S29, Table S8). This is consistent with balancing selection within the static eQTL; however, intermediate minor allele frequencies can also be a signature of soft sweeps (Messer and Petrov 2013). Because static eQTL tend to have higher effect sizes than site specific eQTL (Fig. S31), we re-ran the tests only including eQTL in the top quartile of effect sizes for each population and found the same pattern. GxE eQTL were mostly trans (Fig. 3a, 3b) and had overall lower minor allele frequencies, consistent with purifying selection (Fig. S32, Table S8).

### eQTL regulate expression of candidate genes for leaf phosphorus concentration

Motivated by evidence of GxE interactions in switchgrass leaf P concentration (Zhang et al. 2021), and GxE for expression of phosphorus-related genes in leaves (Fig. 1), we further explored the genetic basis of leaf P variation in the diversity panel. We uncovered GxE for leaf P concentration structured by population: on average, Midwest genotypes had lower phosphorus concentration in Michigan and higher leaf P in Texas, while Gulf and Atlantic populations have higher leaf P in Michigan and lower leaf P in Texas (Fig. 5a).

**Figure 5.**
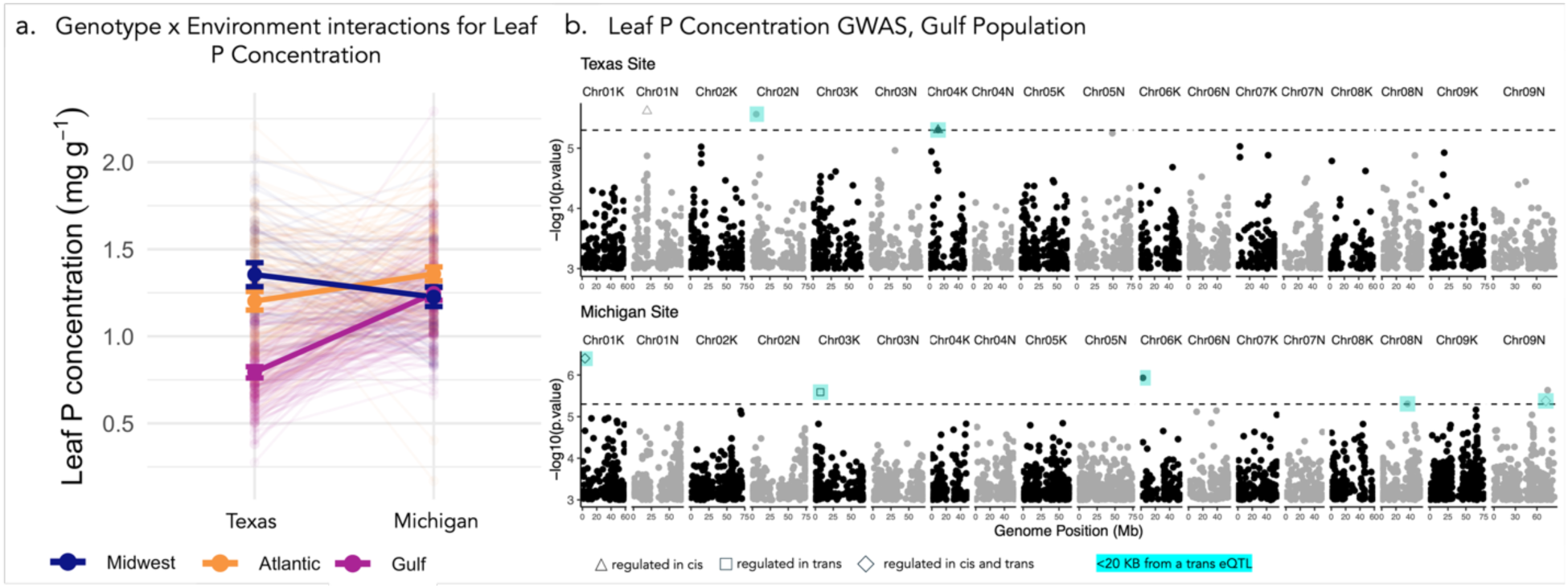
Genotype x environment (GxE) interactions and GWAS of leaf phosphorus (P) concentration in switchgrass. a) GxE interactions for leaf P concentration. Bold lines show the median and interquartile range for leaf P concentration in each sub-population in Texas and Michigan. Pale lines show the P concentration in Texas and Michigan for all individuals. b) Switchgrass leaf P concentration GWAS in the Atlantic population at the Texas and Michigan sites. Triangle points are those whose candidate genes are regulated in cis by an eQTL from Fig. 3; square points are regulated in trans; diamond points are regulated in cis and in trans. Points highlighted in turquoise are candidate SNPs within 20Kb of a trans eQTL from Figure 2.

To understand the molecular underpinnings of variation in leaf P concentration, we conducted a GWAS in the Gulf, Midwest, and Atlantic populations, and identified candidate genes within 20Kb of candidate SNPs. We found 5 Atlantic candidate genes near 4 candidate SNPs (3 in MI, 2 in TX, Fig. 5b), 13 Gulf candidate genes near 9 candidate SNPs (10 in MI, 3 in TX, Fig. S31, Table S9), and 0 Midwest candidate genes. Consistent with the observed GxE for leaf P (Fig. 5a), distinct candidate SNPs and candidate genes were associated with leaf P in Michigan vs. Texas in both the Atlantic and Gulf subpopulations.

Because the genes with expression GxE identified in Fig. 1 were enriched for GO terms related to phosphorus metabolism, we tested the relationship between expression of these genes and leaf phosphorus concentration. After correcting for population structure and false discovery rate, we found that expression of one gene, Pavir.5KG551100 significantly predicts leaf P concentration in the Gulf population at the Texas site (adjusted p-value < 0.018, adjusted R^2^ = 0.141, Fig. S34). This candidate gene has significant GxE for gene expression in AP13 and DAC6 in MI and TX (Fig. 1a) and is within the gene module with the strongest GxE for module eigengene expression (Fig. 1c). Pavir.5KG551100 is a phosphenolpyruvate carboxylase (PEPC), a plant enzyme that is important for C4 photosynthetic carbon assimilation (Carvalho et al. 2024).

Overlaying the leaf P concentration GWAS with the eQTL map and previous leaf P concentration QTL reveals potential mechanistic links between SNP and trait. For example, one of the Gulf candidate genes in Michigan, Pavir.1KG046100, is within a previously identified quantitative trait locus for leaf P in a 4-way mapping population of upland and lowland switchgrass (Zhang et al. 2021). Pavir.1KG046100’s function is associated with DNA binding (Table S9). The expression of Gulf leaf P candidate genes was regulated in cis and in trans by eQTL. 5 of the 9 Gulf candidate SNPs were regulated by eQTL (Fig 5b, Table S9). However, none of the Atlantic candidate genes were regulated by eQTL (Fig. S33). Because candidate SNPs may themselves regulate the expression of genes in trans, or be in linkage disequilibrium with trans eQTL, we investigated whether any candidate SNPs were within 20Kb of trans eQTL and checked whether candidate SNPs were near any eQTL hotspots. 6 of the 9 Gulf candidate genes were within 20KB of a trans eQTL, and one Gulf SNP was near a trans eQTL hotspot that regulates the expression of 41 genes. The complete list of genes regulated by this trans eQTL hotspot, and their annotations and GO terms, is found in Table S10. That candidate SNP is 4Kb downstream of Pavir.6KG022900, a protein phosphatase 2A gene (Fig. 5b, Table S10). 2 of the Atlantic candidate genes are within 20KB of a trans eQTL (Fig. S33, Table S9).

## Discussion

Here, we have characterized the effects of genotype and environment on gene expression and gene regulatory variation in switchgrass. We discovered that differentially expressed genes and gene modules with significant GxE interactions are enriched for genes related to phosphorus metabolism. Through eQTL mapping in the switchgrass diversity panel, we identified thousands of cis and trans eQTL, both static (eQTL mapped in both MI and TX), and dynamic (site-specific or GxE eQTL). In this system, cis eQTL are more consistent across space than trans eQTL which tend to be more dynamic (Fig. 3). Trans eQTL exhibit signatures of stronger purifying selection than cis eQTL, and static eQTL exhibit potential signatures of balancing selection. After discovering that DE genes and GxE gene modules were enriched for processes involving phosphorus, we conducted a genome-wide association study of leaf P concentration in the switchgrass diversity panel and identified candidate genes. To better understand how these candidate genes may influence gene expression, we overlaid our leaf P candidate genes with the cis and trans eQTL identified in the diversity panel. Taken together, this work sheds light on both the molecular underpinnings and evolutionary history of GxE interactions in plants.

### Phosphorus metabolism-related genes exhibit expression GxE

The Texas and Michigan common gardens differ along several key environmental axes, and the differentially expressed genes reflect AP13 and DAC6’s response to that variation. Because the switchgrass genotypes are integrating multiple environmental cues at once, it is difficult to tease apart which specific environmental variable is cueing expression of a particular gene. However, examining enriched GO terms in differentially expressed genes and gene modules can suggest which molecular processes are affected by this variation in climate, and provide future avenues for functional investigation. Acid phosphatases and phosphorus metabolism-related GO terms were overrepresented in the GxE DE genes and the gene module with the strongest GxE interaction (Fig. 1). Acid phosphatase gene expression is a universal response to phosphorus deficiency in plants and other stress conditions (Duff et al. 1991; Duff et al. 1994; Pozo et al. 1999; Baldwin et al. 2001; Vance et al. 2003). The most extreme GxE DE gene, Pavir.4NG062966, is a myo-inositol oxygenase (Table S1). Myo-inositol oxygenase mediates the production of phytic acid, the key storage molecule for phosphorus in plants (Siddique et al. 2014; Kurita et al. 2017; Poirier et al. 2022). Furthermore, in independent *Arabidopsis* and bacterial GWAS, myo-inositol impacts the colonization of phosphate-solubilizing bacteria (O’Banion et al. 2023).

In the common gardens, the Michigan site has higher soil phosphorus than the Texas site, although in absolute terms phosphorus is low in both gardens (Edwards et al. 2023). If there was a simple linear relationship between soil phosphorus levels and acid phosphatase expression, we would expect both genotypes to exhibit higher expression of acid phosphatases in TX and lower expression in MI. Instead, we find a crossed reaction norm, where AP13 acid phosphatase gene expression is lower in Michigan and higher in Texas, and the opposite trend in DAC6 (Fig. 1c, Table S1). This suggests that both genotype and environmental variables interact to shape acid phosphatase expression. Differences in gene expression between upland and lowland switchgrass reflect the integration of diverse environmental signals from the Michigan and Texas sites. For example, the root microbiome of switchgrass exhibits variation based on site and genotype (Edwards et al. 2023). Since microbes play an essential role in converting phosphorus into its bioavailable form (Pi), the microbial community is likely a key variable shaping GxE for expression of phosphorus-related genes. In the diversity panel, we found similar GxE interaction for leaf P concentration at the population level, with a crossed reaction norm between upland and lowland individuals (Fig. 5a). Overall, diversity panel genotypes maintain higher leaf P concentration when grown in a site farther from their population’s region of origin (Fig. 5a). This could indicate that greater stress in a distant environment favors more acquisitive, nutrient-rich leaves (Donovan et al. 2011). In future, controlled experiments are necessary to isolate the specific environmental variables that cause these differences in leaf P concentration and phosphorus-related gene expression between sites.

### eQTL mapping in contrasting sites reveals the evolutionary history of regulatory variation

Mapping regulatory variation across diverse populations and distinct environments yields insights into how selection shapes the regulation of gene expression. In the diversity panel, 19.1 – 45.5% of genes were regulated in cis across populations and sites. Through eQTL mapping, we found substantial trans regulatory variation. We also identified site-specific trans regulatory hotspots in which a single variant regulates the expression of > 50 genes.

Mapping eQTL in contrasting common gardens allows us to assess gene regulatory variation in a context similar to the natural environment in which selection acts on traits. We found that cis eQTL are more likely to be mapped in both sites than trans eQTL, which were more site-specific (Fig. 3). This aligns with previous evidence from experiments mapping eQTL in contrasting conditions, in which cis eQTL were more stable and trans eQTL were more likely to be condition-dependent (Grundberg et al. 2012; Ackermann et al. 2013; Cubillos et al. 2014; Snoek et al. 2017; Lovell et al. 2018; Verta and Jones 2019; Ballinger et al. 2023). We mapped 10 - 15x as many GxE trans eQTL as GxE cis eQTL (Fig. 3b), providing further support that trans eQTL have more dynamic effects across space. All trans regulatory hotspots – trans eQTL that affected the expression of > 50 genes – were site-specific (Fig. 2a). These site-specific regulatory hotspots may act as ‘master switches’ allowing plants to respond to diverse environmental conditions.

Examining the site frequency spectrum reveals information about the evolutionary history of distinct classes of regulatory variants. In our eQTL maps, lower minor allele frequencies in the trans eQTL suggest stronger purifying selection on trans eQTL than cis eQTL. This is consistent with theoretical predictions that mutations affecting gene regulation in trans are more likely to be deleterious because they can alter the expression of many genes simultaneously (McGuigan et al. 2014). Our work contributes to a growing body of empirical evidence that trans eQTL are under purifying selection (Josephs et al. 2020; Brown and Kelly 2022), and extends the inference to show that this evidence of purifying selection is present in static, site-specific, and GxE trans eQTL (Fig. 5, Figs. S27, S28, S30).

We discovered that static eQTL tend to have more intermediate minor allele frequencies, compared to site-specific eQTL (Fig. 5b, Fig. S28-S29, Table S6), which is consistent with balancing selection (Kreitman and Rienzo 2004). Why would selection act to maintain genetic diversity in eQTL with static effects on expression across environments? One possibility is that spatially varying selection has occurred, such that local adaptation has involved purifying selection at narrow spatial scales but balancing across broader spatial scales (Delph and Kelly 2014). However, intermediate allele frequencies may also indicate soft sweeps (Messer and Petrov 2013). Further population genetic analyses are necessary to distinguish the evolutionary history of static and site-specific eQTL. Whether they are static or site-specific, the maintenance of diversity in regulatory variants may be beneficial as the climate changes, as these alleles may facilitate rapid adaptation to climate fluctuations (e.g. Campbell-Staton et al. 2021; Hamann et al. 2021; Ballinger et al. 2023).

### Candidate genes for leaf phosphorus variation are regulated by cis and trans variants

Identifying the genes governing important traits is a fundamental goal of evolutionary genetics. Integrating GWAS with eQTL studies can get us closer to the mechanisms through which genetic variants give rise to traits. Furthermore, conducting GWAS and eQTL mapping in multiple sites allows us to identify specific candidate variants associated with GxE interactions.

Consistent with observed differences in leaf P concentration in Texas and Michigan, we mapped unique candidate genes associated with leaf P at each site (Fig 6b, Fig, S31-32, Table S10). The GWAS revealed a moderately polygenic basis for leaf P concentration, 4 SNPs in the Atlantic population and 9 candidate SNPs in the Gulf population (Fig. 6, Fig. S33). One leaf P concentration candidate gene, Pavir.6KG022900, is a protein phosphatase 2A gene and is likely directly involved in phosphorus metabolism (Huang et al. 2012). Overall, leaf P candidate genes across populations and sites had diverse predicted functions from plant defenses to metabolism, to regulation of gene expression (Table S10). Indeed, 6 of the 9 Gulf candidate SNPs were within 20KB from trans eQTL (Fig. 5b), and regulation of gene expression likely mediates GxE interactions for phosphorus concentration in leaves.

### Conclusions

Genetic diversity and environmental cues interact to shape gene expression and plant traits in nature. To understand the molecular basis and evolutionary history of this phenomenon, it is necessary to conduct experiments that encompass both genetic and environmental variation. Here, we have characterized GxE interactions for traits, gene expression, and gene regulatory variation in a diverse perennial grass. Through eQTL mapping, we discovered that cis eQTL have more consistent effects across environments. Furthermore, we found minor allele frequencies consistent with purifying selection on trans eQTL, and intermediate minor allele frequencies in static eQTL that are consistent with potential balancing selection. Our GWAS concentration revealed GxE interactions and a polygenic basis for variation in leaf P concentration. Taken together, our study thoroughly explores the molecular basis and evolutionary history of variation in gene expression and gene regulatory variation.

## Materials and Methods

### Investigating differential gene expression across contrasting sites in upland and lowland switchgrass

To investigate switchgrass trait and gene expression variation between genotypes and environments, upland (DAC6, Michigan n = 3, Texas n = 3), lowland (AP13, Michigan n = 5, Texas n = 5) parents and AP13 x DAC6 F_1_’s (AP13xDAC6, Texas n = 5, Michigan n = 5) were clonally propagated and reared in a northern environment upland environment at Kellogg Biological Station in Hickory Corners, Michigan (KBSM), and a southern environment at the J.J. Pickle Research Campus in Austin, Texas (PKLE). We measured specific leaf area in all four lines, and collected RNA from AP13, DAC6, and the AP13xDAC6 F_1_’s. Leaves were collected from each plant during the post-flowering stage on May in Texas and June in Michigan. Tissue was flash frozen in liquid nitrogen and RNA was extracted and converted to cDNA using a sigma kit (Merck KGaA, Darmstadt, Germany). cDNA was sequenced on an Illumina NovaSeq 6000 at the Joint Genome Institute.

For the analysis of the AP13 and DAC6 RNAs, we minimized reference bias by generating an AP13 – DAC6 pseudoreference using the AP13v6H1 and whole genome sequence from DAC6 (SRR3709212). DAC6 was trimmed with Trimmomatic (Bolger et al. 2014) and aligned to AP13v6H1 using bwa mem (Li and Durbin 2009) with default parameters; variants were called with GATK_4.3 (McKenna et al. 2010) and filtered to retain biallelic sites with a minimum quality of 29 and a minimum depth of 5. We generated the pseudoreference with the GATK command fastaAlternateReferenceMaker and default parameters. AP13 and DAC6 RNA reads were aligned to the pseudoreference with STAR (Dobin et al. 2013), only retaining reads that mapped to a maximum of two places on the genome; multi-mapping reads were randomly assigned to one location on the genome. Reads were quantified with HTSeq2.0.7 (Anders et al. 2015) with the ‘union’ and ‘stranded=reverse’ parameters. Code for these and subsequent gene expression analyses can be found at github.com/bergcolette/switchgrass_eQTL.

To identify genes with differential expression between AP13 and DAC6 in Michigan and Texas common gardens, we used a linear model accounting for the effects of genotype, site, and genotype x site interactions using DESeq2 (Love et al. 2014). 32269 genes (46% of all AP13v6H1 genes) were retained for analysis after filtering by median expression > 3 and presence in at least 10 individuals. We examined the annotations of the top 10 most significant differentially expressed genes and conducted gene ontology enrichment analyses of the DE genes using the R package clusterProfiler (Yu et al. 2012) for both the GxE DE genes (shaped by site, genotype, and their interaction), and specific contrasts (AP13 vs. DAC6; Michigan vs. Texas).

We next identified modules of co-expressed genes using the package WGCNA (Langfelder and Horvath 2008) with a soft threshold of 5, a merge cut height of 0.3, and a minimum module size of 100 genes. Only genes with a median expression level > 3 were included in the co-expression analysis, and genes were normalized using the variance stabilizing transformation in DESeq (Love et al. 2014). We extracted the top 10 hub genes from each module to identify putative genetic pathways involved in each group of co-expressed genes. To understand the biological processes associated with each module, we conducted gene ontology analysis with the clusterProfiler package in R (Yu et al. 2012) and examined enriched GO terms in each module. We also explored cis regulation of genes through allele-specific expression in AP13xDAC6 F1’s; detailed information and results are available in the Supplementary Material.

### Expression GWAS across diverse switchgrass in upland and lowland environments

To investigate cis and trans regulation of gene expression across diverse switchgrass, we conducted an expression genome wide association study (eGWAS) with the diversity panel presented in Lovell et al. 2021. Whole genome sequence reads were trimmed with Trimmomatic (Bolger et al. 2014) and aligned to the AP13v6H1 reference genome using bwa mem (Li and Durbin 2009) with default parameters. Genotypes were called using GATK_4.2 (McKenna et al. 2010) and only variant sites were retained. The resulting vcf was filtered to only retain SNPs with less than 1% of sites missing, minimum depth of 10, minor allele frequency > 0.05, and minimum quality of 29. We split the diversity panel into populations identified by Lovell et al. 2021: Midwest, Gulf, and Atlantic, and LD-pruned the split variant files using the command indep-pairwise in plink2 (Purcell et al. 2007) with a window size of 5000 bp, a step size of 50 bp, and an LD threshold of 0.5. Missing sites in the LD-filtered files were imputed with beagle (Browning and Browning 2016). Because highly heterozygous sites and very rare alleles can lead to Type I errors in eGWAS (Josephs et al. 2015), we filtered the sites to those with and a heterozygous genotype frequency < 0.7. and a minor allele of frequency > 0.05.

RNA for eQTL mapping was extracted from the same individuals in the whole-genome sequenced diversity panel. These expression data were first presented in Izquierdo et al. 2026, and more details on the molecular work can be found there. In brief, tissue was collected from post-flowering individuals May 5^th^-6^th^ 2021 in Texas and June 22^nd^-24^th^ in Michigan between 10AM and noon. Leaf tissue was flash frozen in liquid nitrogen within less than a minute of removal from the plant and RNA was extracted using a Direct-zol™ RNA Miniprep Kit (Zymo Research). RNAseq was performed on an Illumina NovaSeq 6000 at the Joint Genome Institute. RNA reads were aligned to the AP13v6H1 reference genome with HISAT2_2.2.0, and featureCounts (Liao et al. 2014) was used to produce raw read counts.

Reads were filtered to only retain genes with a median read count > 3 across all individuals. We scaled samples by library size by dividing raw read counts by the total number of reads for that sample and normalized these proportions and taking the square root of the value. To map differences in gene expression between sites, we subtracted the size-standardized square root transformed expression in Michigan from that in Texas at each gene. Because most genotypes in the diversity panel do not have replicates, we were unable to account for variance in gene expression within each genotype and therefore acknowledge that the difference in gene expression between sites may contain some statistical noise.

Because the switchgrass diversity panel has strong population structure (Lovell et al. 2021) we split the diversity panel into subpopulations: Midwest (n = 86), Gulf (n = 171), and Atlantic (n = 190) for all eGWAS analyses. Cis and trans regulatory SNPs were mapped with the package matrixEQTL (Shabalin 2012). For each population, we conducted three eQTL scans: one scan mapping the effects of genotype on expression in Texas, one mapping the effects of genotype in Michigan, and another where we mapped the effects of genotype on the difference in expression between Michigan and Texas. We defined cis regulatory SNPs as those that affect expression of a gene < 1Mb away from the variant, and trans regulatory SNPs were defined as variants that affect expression of a gene > 1Mb away. Within each population, we controlled for relatedness by performing principal components analysis of each vcf in plink (Purcell et al., 2007) and including the first five PC’s as covariates.

For all eQTL scans, we only retained eQTL with a minor allele frequency > 0.05 and a false discovery rate < 0.05. In all eGWAS maps, we identified genes regulated by with an implausible number (n > 100) of trans eQTL evenly distributed across the genome (423 genes in Gulf, 81 genes in Atlantic, 10 genes in Midwest). These genes were more likely to exhibit a non-normal distribution of gene expression even after size standardization and square-root transformation (Fig. S35). These results may be attributed in part to the transformed gene expression distributions violating the assumptions of matrixEQTL’s linear model: removing the genes whose expression significantly deviated from normality (p value of Shapiro-wilk test < 0.05) left only 29 genes regulated by > 100 eQTL across all populations. However, this filtering by non-normality would remove over half of all analyzed genes, most of which are not associated with unusually many eQTL. Thus, while we acknowledge that any normalization method will influence eQTL results (see Brown and Kelly 2022), we assert that it is more conservative to retain all genes, even those whose expression is associated with 100s of eQTL, in subsequent analyses. We also discovered trans eQTL that appeared as a diagonal band on a homeologous chromosome (i.e., a diagonal band of trans eQTL on Chr09K regulating genes on Chr09N, Fig. 2a, Fig. S19-S20). We hypothesize that these are the result of mis-mapping, in which reads are erroneously mapped to the corresponding gene on the homeologous chromosome, but retain them in further analyses.

### Investigating the evolutionary history of regulatory variants

We examined the mapped eQTL in Texas and Michigan and divided the eQTL into those that were only mapped in one site (site-specific), and those that were mapped in both sites. We categorized eQTL that significantly predicted a change in gene expression between TX and MI as GxE eQTL. To understand whether cis or trans eQTL tend to have stronger effects on gene expression, we compared the effect sizes of cis and trans eQTL in all populations and sites using t-tests.

For cis and trans eQTL that were mapped in both sites, we characterized how well a variant’s effect on expression in Michigan predict its effect in Texas using a linear regression in R. To understand whether cis or trans eQTL were more consistent across space, we compared the proportion of static and site-specific cis and trans eQTL in each population using chi-square tests. We also compared absolute counts of GxE cis and trans eQTL.

To investigate the evolutionary history of cis and trans eQTL, we compared the minor allele frequency of cis eQTL, trans eQTL, and all SNPs for each population and site with chi-square-tests. Similarly, we compared minor allele frequencies in static and site-specific eQTL. Furthermore, we used linear regression to test for the relationship between absolute eQTL effect size and minor allele frequency. We found a negative relationship between minor allele frequency and affect size and conducted further testing to verify whether the difference in MAF between trans and cis eQTL could be attributed to differences in effect sizes. Therefore, we re-ran our chi-square tests but only included cis and trans eQTL whose effect sizes were in the top quartile.

### Genome wide association study of leaf phosphorus concentration

To identify genomic regions associated with leaf phosphorus (P) concentration in switchgrass, we conducted a genome wide association study (GWAS) with the same diversity panel used in our eGWAS and presented in Lovell et al (2021). For each site, leaves were collected for every individual plant and oven-dried it at 60 °C for > 72 h. The dried leaf material was then ground using plastic vials and yttria-stabilized zirconium beads, the leaf P concentration was analyzed through inductively coupled plasma mass spectrometry (ICP-MS; SCIEX Elan 6000 DRC-e mass spectrometer, Perkin-Elmer, Sheldon, CT) connected to a PFA microflow nebulizer (Elemental Scientific, Omaha, NE) and an Apex HF desolvator (Elemental Scientific, Omaha, NE).

To calculate site heritability for leaf-phosphorus concentration associated SNPs we followed a two-step method as previously described in (George et al. 2000). Variance component analysis was performed with ASReml (VSNi), using a genomic relationship matrix calculated through the van Raden method (VanRaden 2008) for the full diversity panel. The kinship matrix, based on SNPs called against the P. virgatum AP13v6H1 reference genome (Izquierdo et al 2026), was calculated and used for the site-specific heritability mapping using a mixed-effect model in the form of:

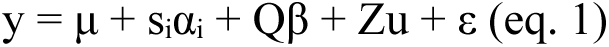

in which y represents the vector of observed total leaf phosphorus concentrations, μ is the intercept, sᵢ is the genotype vector for the i-th SNP being tested, αᵢ is the fixed additive effect of the i-th SNP, Q represents population structure covariates (with three principal components), β is the vector of regression coefficients associated with population structure, Z is the incidence matrix linking individuals to their genomic effects, u is random polygenic genetic effects, and ε is the residual error vector. The variance assumptions remain unchanged as:

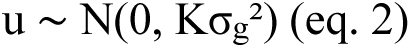

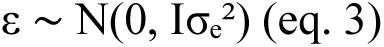

where K is the genomic relationship matrix, σ_g_² is the polygenic variance, and σₑ² is the residual variance. It is important to note that, although Q represents the population structure covariate, and the models were implemented with three principal components (coincidently matching the number of subpopulations), the GWAS pipeline was applied to each subpopulation and site combination separately. Prior to GWAS, marker filtering was performed with a minor allele frequency (MAF) threshold set to 0.05 and a heterozygosity parameter to 0.95. The association significance threshold was set to p < 5 × 10⁻⁴. All GWAS data processing, statistical analyses, and visualization procedures were conducted in R using the ASReml, bigsnpr, data.table, dplyr, and ggplot2 packages.

Next, we investigated whether the expression genes identified in the differential expression and gene module analyses is associated with leaf P concentration in the diversity panel. We ran linear models with leaf P concentration as the response variable and normalized gene expression, with the first five principal components of the genotype matrix included as fixed effects to account for population structure, using the function lm() in R. We ran separate linear models for each of the significant GxE DE genes (Fig. 1a, Table S1), the genes with significant differential expression between site and genotype but no GxE (Table S2), and all the genes within the tan module, which had the strongest GxE for module eigengene expression (Figure 1c). We corrected for false discovery rate by calculating an adjusted p-value with a Benjamini-Hochberg procedure with the R function p.adjust().

We overlaid the leaf P concentration candidate SNPs from the GWAS with the eQTL mapping data from each population and site. We characterized whether candidate genes were regulated in cis or in trans in the eQTL maps and identified candidate SNPs which were <20Kb from a trans eQTL, taking note of whether candidate SNPs that were close to trans eQTL hotspots. After identifying a candidate SNP near an eQTL hotspot, we examined the annotations and associated GO terms of the genes regulated by that variant and described their potential involvement in leaf P concentration. We used the switchgrass AP13v6H1 annotation to characterize the potential mechanisms through which these candidate SNPs influence leaf P concentration.

## Acknowledgements

We gratefully acknowledge Dr. Emily Josephs and Dr. Sarah Lebeis for their helpful comments on the analyses and manuscript. We thank the many technicians and volunteers who helped to maintain these field experiments over the years and collected the tissue for this study. This research was supported by the DOE Office of Science, Office of Biological and Environmental Research (BER), grant no. DE-SC0021126 and DE-SC0026058. This material is based upon work supported by the Great Lakes Bioenergy Research Center, U.S. Department of Energy, Office of Science, Biological and Environmental Research Program under Award Number DE-SC0018409. Support for this research was also provided by the NSF Long-term Ecological Research Program (DEB-2224712) at the Kellogg Biological Station and by Michigan State University AgBioResearch. The work (proposal: 10.46936/10.25585/60000516) conducted by the U.S. Department of Energy Joint Genome Institute (https://ror.org/04xm1d337), a DOE Office of Science User Facility, is supported by the Office of Science of the U.S. Department of Energy operated under Contract No. DE-AC02-05CH11231.

## Author Contributions

C.S. Berg – Conceptualization, Data Curation, Formal Analysis, Original Draft Preparation. C.G. Pereira – Investigation, Formal Analysis, Review and Editing, A.S Sreedasyam – Formal Analysis, J. Lovell – Formal Analysis, X. Weng – Investigation, J. Bonette – Investigation, Data Curation. Y. Yoshinaga – Formal Analysis, C. Daum – Formal Analysis, A. Lipzen – Formal Analysis, J. Schmutz – Formal Analysis, T. Juenger – Conceptualization, Supervision, Resources, Review and Editing. D.B. Lowry – Conceptualization, Supervision, Resources, Review and Editing

## Data Availability

The raw RNA sequencing data for AP13, DAC6, and F1’s are available under PRJNA440167 at https://www.ncbi.nlm.nih.gov/bioproject/PRJNA440167. The raw whole genome sequencing data for the diversity panel are available under BioProject PRJNA265643 at https://www.ncbi.nlm.nih.gov/bioproject/PRJNA265643. The raw RNA sequencing data for the diversity panel are available under BioProject PRJNA1402996 at https://www.ncbi.nlm.nih.gov/bioproject/PRJNA1402996.

