## Supplemental Text, Figures, and Tables S3 and S5-S8 for "Genotype and environment shape gene regulatory variation in switchgrass"

Supplementary Text

### Gene ontology analyses of differentially expressed genes and gene modules

We have provided figures from the GO term enrichment analysis from weighted gene coexpression network analyses (Figs S1 – S13) for all modules with more than one significantly enriched GO term. We do not have figures for the light cyan module, which was only enriched for lipid binding, the green module, which was only enriched for intracellular protein transport, the black module, which was only enriched for vesicle mediated transport, and the pink, cyan, and black modules, which were not significantly enriched for any GO terms.

### Mapping allele-specific expression

We quantified cis-regulatory effects by measuring allele-specific expression (ASE) in AP13 × DAC6 $F_1$ hybrids grown in the Michigan and Texas common gardens. First, we identified one-to-one orthologous genes between AP13 and DAC6 using OrthoFinder v2.5.5 (Emms & Kelly, 2019). RNA-seq reads were then competitively aligned against the AP13v6H1 and DAC6v6H1 reference genomes using STAR v2.7.11b (Dobin *et al.*, 2013). Reads were classified as either uniquely mapped (mapping preferentially to one of the two reference genomes) or shared (mapping equally well to both genomes). For each orthologous gene, allele-specific read counts were calculated by summing the uniquely mapped reads and assigning half of the shared reads to each allele. This strategy accounts for ambiguously mapped reads while reducing reference-mapping bias and enabling balanced quantification of expression from the two parental alleles. To calculate the degree of allele-specific expression, we retained genes overlapping with expected heterozygous sites (i.e., sites that are fixed between the AP13 and DAC6 parents) and filtered to only retain genes with a total depth of 30, that were present in at least 3 of the 5 $F_1$ replicates in each site. We assessed deviation from the expected allelic ration of 0.5 with binomial tests. After correcting for multiple testing, we considered genes to have ASE if the p-value of the binomial test was <2.2e-6. To assess whether ASE was consistent across sites, we quantified the overlap in ASE genes in both Michigan and Texas.

A substantial proportion of genes in AP13xDAC6 $F_1$'s exhibited allele-specific expression, and the proportion of bias in each direction was similar in the two sites. In the AP13xDAC6 $F_1$'s at the Michigan site, 14645 of the 22642 one-to-one orthologous genes with fixed SNPs between parents exhibited allele-specific expression, with 7574 genes biased toward the AP13 allele and 7071 genes biased toward the DAC6 allele (Fig. S27). ASE patterns were similar at the Texas site: of 15082 ASE genes, 7830 were biased toward the AP13 allele and 7252 were biased toward the DAC6 allele (Fig. S27). Most of the allele-specific expression was consistent across sites: 13230 ASE genes (90.3% of MI ASE genes, 87.7% of TX ASE genes) had expression bias in the same direction in both Michigan and Texas (Fig S27). We identified 1307 MI-specific ASE genes and 1744 TX-specific ASE genes. Furthermore, only 108 significant ASE genes had bias in the opposite direction between sites (Fig. S27).

Across sites and populations, some of the cis-regulated genes from the QTL map overlapped with the cis-regulated genes identified through ASE mapping (Fig. 2, Table S7). However, there was not significant enrichment for overlap between the ASE cis-regulated genes and the eQTL cis-regulated genes. This suggests unique divergence between AP13 and DAC6 that is not fully captured in the diversity panel and differences in power between the two analyses.

#### Examining integration of gene regulation within and between subgenomes

Switchgrass is an allotetraploid with disomic inheritance (Okada *et al.*, 2010; Triplett *et al.*, 2012). Because polyploids can maintain distinct co-expression networks in each subgenome (Adams & Wendel, 2005), we compared the number of eQTL in each subgenome and compared trans-regulation of genes within vs. between each subgenome. We found 2 – 7% more trans eQTL regulating gene expression within their subgenomes vs. between subgenomes (Fig. S22) in all three subpopulations. However, we did not find consistent differences in the number of eQTL in the dominant (N) vs. subdominant (K) subgenomes: for some chromosomes, the N homeolog harbored more eQTL; for other chromosomes the K homeolog contained more eQTL (Fig. S23). Overall, this suggests high integration of gene expression between subgenomes, and minimal distinction between the K and N subgenomes in terms of number and distribution of eQTL.

### Supplementary Tables

Supplementary Table 1. Genes whose expression was significantly explained by habitat (MI vs. TX), genotype (AP13 vs. DAC6), and their interaction, attached in Bergetal_TableS1.csv. Interaction_type column indicates whether the GxE pattern is rank-changing or variance changing. Zhangetal_2021_QTL_overlap column indicates whether the gene is within the switchgrass leaf phosphorus concentration QTL mapped in Zhang et al. 2021.

Supplementary Table 2. Table S2. Genes with significant differential expression between habitat (MI vs. TX) and genotype (AP13 vs. DAC6), but without a significant GxE interaction, attached in Bergetal_TableS2.csv.

Supplementary Table 3a-o. Model outputs for ANOVAs of the relationship between module eigengene expression and genotype, habitat, and their interaction.

3a. Model output - Tan module eigengene

| Tan ME | Df | Sum Sq | Mean S1 | F value | Pr(>F) | significance |
|---|---|---|---|---|---|---|
| habitat | 1 | 0.0942 | 0.0942 | 12.356 | 0.00656 | ** |
| genotype | 1 | 0.0711 | 0.0711 | 9.334 | 1.37E-02 | * |
| habitat:genotype | 1 | 0.5441 | 0.5441 | 71.388 | 1.43E-05 | *** |
| Residuals | 9 | 0.0686 | 0.0076 | | | |

3b. Model output = Magenta module eigengene

| Magenta ME | Df | Sum Sq | Mean S1 | F value | Pr(>F) | significance |
|---|---|---|---|---|---|---|
| habitat | 1 | 0.3181 | 0.3181 | 11.225 | 0.00852 | ** |

| | | | | | | |
|---|---|---|---|---|---|---|
| genotype | 1 | 0.0055 | 0.0055 | 0.193 | 0.67079 | |
| habitat:genotype | 1 | 0.4007 | 0.4007 | 14.14 | 0.00448 | ** |
| Residuals | 9 | 0.255 | 0.0283 | | | |

3c. Model output – Turquoise module eigengene

| Turquoise | Df | Sum Sq | Mean S1 | F value | Pr(>F) | significance |
|---|---|---|---|---|---|---|
| habitat | 1 | 0.0132 | 0.0132 | 17.841 | 0.00223 | ** |
| genotype | 1 | 0.9536 | 0.9536 | 1291.384 | 4.95E-11 | *** |
| habitat:genotype | 1 | 0.0027 | 0.0027 | 3.612 | 0.08983 | . |
| Residuals | 9 | 0.0066 | 0.0007 | | | |

3d. Model output – Salmon module eigengene

| Salmon | Df | Sum Sq | Mean S1 | F value | Pr(>F) | significance |
|---|---|---|---|---|---|---|
| habitat | 1 | 0.2814 | 0.28144 | 5.611 | 0.042 | * |
| genotype | 1 | 0.129 | 0.12902 | 2.572 | 0.143 | |
| habitat:genotype | 1 | 0.0168 | 0.01682 | 0.335 | 0.577 | |
| Residuals | 9 | 0.4514 | 0.05016 | | | |

3e. Model output – Purple module eigengene

| Purple | Df | Sum Sq | Mean S1 | F value | Pr(>F) | significance |
|---|---|---|---|---|---|---|
| habitat | 1 | 0.0813 | 0.08126 | 0.967 | 0.351 | |
| genotype | 1 | 0.1064 | 0.10636 | 1.266 | 0.29 | |
| habitat:genotype | 1 | 0.0021 | 0.00207 | 0.025 | 0.879 | |
| Residuals | 9 | 0.7564 | 0.08404 | | | |

3f. Model output – Grey60 module eigengene

| grey60 | Df | Sum Sq | Mean S1 | F value | Pr(>F) | significance |
|---|---|---|---|---|---|---|
| habitat | 1 | 0.19513 | 0.19513 | 9.125 | 0.0145 | * |
| genotype | 1 | 0.08025 | 0.08025 | 3.753 | 0.0847 | . |
| habitat:genotype | 1 | 0.06445 | 0.06445 | 3.014 | 0.1166 | |
| Residuals | 9 | 0.19246 | 0.02138 | | | |

3g. Model output – Green module eigengene

| green | Df | Sum Sq | Mean S1 | F value | Pr(>F) | significance |
|---|---|---|---|---|---|---|
| habitat | 1 | 0.744 | 0.744 | 323.353 | 2.31E-08 | *** |
| genotype | 1 | 0.1187 | 0.1187 | 51.595 | 5.18E-05 | *** |
| habitat:genotype | 1 | 0.0191 | 0.0191 | 8.309 | 0.0181 | * |
| Residuals | 9 | 0.0207 | 0.0023 | | | |

3h. Model output – yellow module eigengene

| yellow | Df | Sum Sq | Mean S1 | F value | Pr(>F) | significance |
|---|---|---|---|---|---|---|
| habitat | 1 | 0.3457 | 0.3457 | 103.638 | 3.08E-06 | *** |
| genotype | 1 | 0.4526 | 0.4526 | 135.69 | 9.91E-07 | *** |
| habitat:genotype | 1 | 0.013 | 0.013 | 3.884 | 0.0803 | . |
| Residuals | 9 | 0.03 | 0.0033 | | | |

3i. Model output – LightCyan module eigengene

| light cyan | Df | Sum Sq | Mean S1 | F value | Pr(>F) | significance |
|---|---|---|---|---|---|---|
| habitat | 1 | 0.2655 | 0.2655 | 9.757 | 0.01225 | * |
| genotype | 1 | 0.0912 | 0.0912 | 3.352 | 0.10038 | |
| habitat:genotype | 1 | 0.3889 | 0.3889 | 14.293 | 0.00434 | ** |
| Residuals | 9 | 0.2449 | 0.0272 | | | |

3j. Model output – Pink module eigengene

| Pink | Df | Sum Sq | Mean S1 | F value | Pr(>F) | significance |
|---|---|---|---|---|---|---|
| habitat | 1 | 0.3808 | 0.3808 | 10.216 | 0.0109 | * |
| genotype | 1 | 0.0487 | 0.0487 | 1.307 | 0.2824 | |
| habitat:genotype | 1 | 0.003 | 0.003 | 0.08 | 0.7832 | |
| Residuals | 9 | 0.3355 | 0.0373 | | | |

3k. Model output – Greenyellow module eigengene

| Greenyellow | Df | Sum Sq | Mean S1 | F value | Pr(>F) | significance |
|---|---|---|---|---|---|---|
| habitat | 1 | 0.081 | 0.081 | 5.566 | 0.042657 | * |
| genotype | 1 | 0.1261 | 0.1261 | 8.67 | 0.016368 | * |
| habitat:genotype | 1 | 0.5848 | 0.5848 | 40.192 | 0.000135 | *** |
| Residuals | 9 | 0.131 | 0.0146 | | | |

3l. Model output – Brown module eigengene

| Brown | Df | Sum Sq | Mean S1 | F value | Pr(>F) | significance |
|---|---|---|---|---|---|---|
| habitat | 1 | 0.3765 | 0.3765 | 124.401 | 1.43E-06 | *** |
| genotype | 1 | 0.4382 | 0.4382 | 144.773 | 7.53E-07 | *** |
| habitat:genotype | 1 | 0.0266 | 0.0266 | 8.781 | 0.0159 | * |
| Residuals | 9 | 0.0272 | 0.003 | | | |

3m. Model output – Black module eigengene

| Black | Df | Sum Sq | Mean S1 | F value | Pr(>F) | significance |
|---|---|---|---|---|---|---|
| habitat | 1 | 0.3542 | 0.3542 | 21.302 | 0.00126 | ** |
| genotype | 1 | 0.0748 | 0.0748 | 4.497 | 0.06298 | . |
| habitat:genotype | 1 | 0.0931 | 0.0931 | 5.598 | 0.04219 | * |

|  |  |  |  |
|---|---|---|---|
| Residuals | 9 | 0.1496 | 0.0166 |

3n. Model output – Red module eigengene

| Red | Df | Sum Sq | Mean S1 | F value | Pr(>F) | significance |
|---|---|---|---|---|---|---|
| habitat | 1 | 0.8094 | 0.8094 | 201.517 | 1.82E-07 | *** |
| genotype | 1 | 0.0555 | 0.0555 | 13.831 | 0.00478 | ** |
| habitat:genotype | 1 | 0.0005 | 0.0005 | 0.124 | 0.73256 |  |
| Residuals | 9 | 0.0361 | 0.004 |  |  |  |

3o. Model output – MidnightBlue module eigengene

| Midnightblue | Df | Sum Sq | Mean S1 | F value | Pr(>F) | significance |
|---|---|---|---|---|---|---|
| habitat | 1 | 0.0796 | 0.07962 | 1.434 | 0.262 |  |
| genotype | 1 | 0.1227 | 0.12269 | 2.21 | 0.171 |  |
| habitat:genotype | 1 | 0.0191 | 0.01914 | 0.345 | 0.571 |  |
| Residuals | 9 | 0.4996 | 0.05551 |  |  |  |

3p. Model output – Cyan module eigengene

| Cyan | Df | Sum Sq | Mean S1 | F value | Pr(>F) | significance |
|---|---|---|---|---|---|---|
| habitat | 1 | 0.122 | 0.122 | 3.619 | 0.08956 | . |
| genotype | 1 | 0.4627 | 0.4627 | 13.725 | 0.00488 | ** |
| habitat:genotype | 1 | 0.0701 | 0.0701 | 2.081 | 0.18306 |  |
| Residuals | 9 | 0.3034 | 0.0337 |  |  |  |

3q. Model output – Blue module eigengene

| Blue | Df | Sum Sq | Mean S1 | F value | Pr(>F) | significance |
|---|---|---|---|---|---|---|
| habitat | 1 | 0.0078 | 0.0078 | 8.78 | 0.0159 | * |
| genotype | 1 | 0.944 | 0.944 | 1056.386 | 1.22E-10 | *** |
| habitat:genotype | 1 | 0 | 0 | 0.003 | 0.9554 |  |
| Residuals | 9 | 0.008 | 0.0009 |  |  |  |

Supplementary Table 4: Top 10 hub genes of all gene coexpression modules in AP13 and DAC6 reared in Michigan and Texas, identified by Weighted Gene Coexpression Network Analysis, attached as Bergetal_TableS4.csv

Supplementary Table 5: Number of cis and trans eQTLs in each population and habitat.

| Population | Type | Site | # eQTL | # genes |
|---|---|---|---|---|
| Midwest | cis | MI_only | 8213 | 5099 |
| Midwest | cis | TX_and_MI | 3715 | 2304 |
| Midwest | cis | TX_only | 3253 | 2603 |
| Midwest | trans | MI_only | 19306 | 6332 |

| | | | | |
|---|---|---|---|---|
| Midwest | trans | TX_and_MI | 2102 | 759 |
| Midwest | trans | TX_only | 16779 | 5608 |
| Midwest | cis | GxE | 26 | 24 |
| Midwest | trans | GxE | 2453 | 1028 |
| Atlantic | cis | MI_only | 14361 | 7656 |
| Atlantic | cis | TX_and_MI | 13246 | 5617 |
| Atlantic | cis | TX_only | 10284 | 6483 |
| Atlantic | trans | MI_only | 38045 | 10822 |
| Atlantic | trans | TX_and_MI | 14904 | 1903 |
| Atlantic | trans | TX_only | 31311 | 9323 |
| Atlantic | cis | GxE | 54 | 48 |
| Atlantic | trans | GxE | 584 | 365 |
| Gulf | cis | MI_only | 18384 | 10131 |
| Gulf | cis | TX_and_MI | 14802 | 6117 |
| Gulf | cis | TX_only | 23715 | 12388 |
| Gulf | trans | MI_only | 121410 | 20108 |
| Gulf | trans | TX_and_MI | 29941 | 2867 |
| Gulf | trans | TX_only | 178299 | 25965 |
| Gulf | cis | GxE | 34 | 29 |
| Gulf | trans | GxE | 6928 | 2245 |

Supplementary Table 6. eQTL hotspots. Chromosome and bp position of trans eQTL that regulate the expression of > 50 genes.

| genome_Chrom | genome_bp | presence | type | # genes regulated | population |
|---|---|---|---|---|---|
| Chr01K | 32301507 | TX_only | trans | 52 | Gulf |
| Chr01K | 35377587 | TX_only | trans | 413 | Gulf |
| Chr01K | 53128033 | TX_only | trans | 81 | Gulf |
| Chr01K | 56721489 | MI_only | trans | 56 | Gulf |
| Chr01K | 56895910 | TX_only | trans | 150 | Gulf |
| Chr01K | 57315501 | TX_only | trans | 101 | Gulf |
| Chr01K | 57424127 | TX_only | trans | 118 | Gulf |
| Chr01N | 1874991 | TX_only | trans | 85 | Gulf |
| Chr01N | 21298272 | MI_only | trans | 78 | Gulf |
| Chr01N | 53613853 | TX_only | trans | 398 | Gulf |
| Chr01N | 54454683 | TX_only | trans | 57 | Gulf |
| Chr01N | 57440699 | MI_only | trans | 71 | Gulf |
| Chr01N | 66333130 | TX_only | trans | 101 | Gulf |
| Chr02K | 53134 | MI_only | trans | 67 | Gulf |

| | | | | | |
|---|---|---|---|---|---|
| Chr02K | 73954 | MI_only | trans | 130 | Gulf |
| Chr02K | 84726 | MI_only | trans | 120 | Gulf |
| Chr02K | 5102789 | TX_only | trans | 78 | Gulf |
| Chr02K | 40171713 | TX_only | trans | 338 | Gulf |
| Chr02K | 51747114 | TX_only | trans | 68 | Gulf |
| Chr02K | 65052737 | MI_only | trans | 62 | Gulf |
| Chr02N | 13768330 | TX_only | trans | 73 | Gulf |
| Chr02N | 17077748 | TX_only | trans | 78 | Gulf |
| Chr02N | 23522775 | MI_only | trans | 54 | Gulf |
| Chr02N | 62020397 | MI_only | trans | 61 | Gulf |
| Chr02N | 65366104 | TX_only | trans | 65 | Gulf |
| Chr02N | 69597293 | TX_only | trans | 128 | Gulf |
| Chr02N | 70565539 | TX_only | trans | 60 | Gulf |
| Chr02N | 73201432 | MI_only | trans | 53 | Gulf |
| Chr02N | 73281882 | MI_only | trans | 52 | Gulf |
| Chr02N | 73543664 | MI_only | trans | 63 | Gulf |
| Chr03K | 11104266 | TX_only | trans | 267 | Gulf |
| Chr03K | 17337330 | TX_only | trans | 144 | Gulf |
| Chr03K | 29708527 | TX_only | trans | 217 | Gulf |
| Chr03K | 46918706 | MI_only | trans | 51 | Gulf |
| Chr03K | 59702834 | TX_only | trans | 71 | Gulf |
| Chr03K | 61484971 | MI_only | trans | 56 | Gulf |
| Chr03N | 1833363 | TX_only | trans | 55 | Gulf |
| Chr03N | 2095815 | TX_only | trans | 84 | Gulf |
| Chr03N | 2118802 | TX_only | trans | 54 | Gulf |
| Chr03N | 3711508 | TX_only | trans | 72 | Gulf |
| Chr03N | 11214412 | TX_only | trans | 80 | Gulf |
| Chr03N | 15182209 | TX_only | trans | 101 | Gulf |
| Chr03N | 19337383 | MI_only | trans | 51 | Gulf |
| Chr04K | 314533 | MI_only | trans | 59 | Gulf |
| Chr04K | 19118307 | TX_only | trans | 72 | Gulf |
| Chr04K | 36706081 | MI_only | trans | 51 | Gulf |
| Chr04K | 38325822 | TX_only | trans | 109 | Gulf |
| Chr04K | 39967129 | TX_only | trans | 57 | Gulf |
| Chr04K | 41373355 | MI_only | trans | 66 | Gulf |
| Chr04N | 1610278 | TX_only | trans | 73 | Gulf |
| Chr04N | 9755725 | MI_only | trans | 60 | Gulf |
| Chr05K | 977179 | TX_only | trans | 104 | Gulf |
| Chr05K | 9959811 | TX_only | trans | 54 | Gulf |

| | | | | | |
|---|---|---|---|---|---|
| Chr05K | 25096708 | TX_only | trans | 60 | Gulf |
| Chr05K | 45336355 | TX_only | trans | 79 | Gulf |
| Chr05K | 50717435 | TX_only | trans | 76 | Gulf |
| Chr05K | 51242549 | TX_only | trans | 66 | Gulf |
| Chr05K | 53075288 | MI_only | trans | 66 | Gulf |
| Chr05K | 58341585 | TX_only | trans | 79 | Gulf |
| Chr05N | 13674092 | TX_only | trans | 56 | Gulf |
| Chr05N | 16566740 | TX_only | trans | 75 | Gulf |
| Chr05N | 18020257 | TX_only | trans | 79 | Gulf |
| Chr05N | 18522806 | TX_only | trans | 59 | Gulf |
| Chr05N | 19471181 | TX_only | trans | 85 | Gulf |
| Chr05N | 27536254 | TX_only | trans | 54 | Gulf |
| Chr05N | 27568593 | TX_only | trans | 61 | Gulf |
| Chr05N | 41979803 | TX_only | trans | 196 | Gulf |
| Chr05N | 48850252 | TX_only | trans | 222 | Gulf |
| Chr05N | 51070900 | MI_only | trans | 58 | Gulf |
| Chr05N | 58454370 | TX_only | trans | 146 | Gulf |
| Chr05N | 58454712 | TX_only | trans | 66 | Gulf |
| Chr05N | 58839161 | TX_only | trans | 89 | Gulf |
| Chr05N | 66880804 | TX_only | trans | 438 | Gulf |
| Chr05N | 72844150 | TX_only | trans | 61 | Gulf |
| Chr05N | 74391122 | MI_only | trans | 56 | Gulf |
| Chr06K | 44545166 | TX_only | trans | 117 | Gulf |
| Chr06N | 29403156 | MI_only | trans | 56 | Gulf |
| Chr06N | 34063848 | MI_only | trans | 62 | Gulf |
| Chr06N | 38214314 | MI_only | trans | 199 | Gulf |
| Chr06N | 53091510 | TX_only | trans | 65 | Gulf |
| Chr07K | 20904809 | TX_only | trans | 56 | Gulf |
| Chr07K | 47962490 | MI_only | trans | 62 | Gulf |
| Chr07N | 26267228 | TX_only | trans | 61 | Gulf |
| Chr07N | 28226985 | TX_only | trans | 59 | Gulf |
| Chr07N | 31895095 | TX_only | trans | 78 | Gulf |
| Chr07N | 37130122 | MI_only | trans | 57 | Gulf |
| Chr07N | 40553750 | TX_only | trans | 88 | Gulf |
| Chr07N | 50175610 | TX_only | trans | 100 | Gulf |
| Chr07N | 51408225 | TX_only | trans | 63 | Gulf |
| Chr08N | 35681378 | MI_only | trans | 114 | Gulf |
| Chr09K | 3282454 | TX_only | trans | 77 | Gulf |
| Chr09K | 3314122 | TX_only | trans | 151 | Gulf |

| | | | | | |
|---|---|---|---|---|---|
| Chr09K | 4831356 | TX_only | trans | 51 | Gulf |
| Chr09K | 13919429 | MI_only | trans | 62 | Gulf |
| Chr09K | 25988439 | TX_only | trans | 51 | Gulf |
| Chr09K | 44390385 | TX_only | trans | 64 | Gulf |
| Chr09K | 51731003 | MI_only | trans | 74 | Gulf |
| Chr09K | 61999986 | TX_only | trans | 85 | Gulf |
| Chr09K | 67017281 | MI_only | trans | 60 | Gulf |
| Chr09N | 6762873 | MI_only | trans | 52 | Gulf |
| Chr09N | 18270074 | MI_only | trans | 63 | Gulf |
| Chr09N | 20498967 | TX_only | trans | 96 | Gulf |
| Chr09N | 38828745 | TX_only | trans | 156 | Gulf |
| Chr09N | 52739731 | TX_only | trans | 370 | Gulf |
| Chr09N | 52922050 | TX_only | trans | 89 | Gulf |
| Chr01K | 9214380 | TX_only | trans | 67 | Midwest |
| Chr01N | 24177862 | TX_only | trans | 103 | Midwest |
| Chr02N | 50382741 | TX_only | trans | 80 | Midwest |
| Chr03K | 17862805 | MI_only | trans | 64 | Midwest |
| Chr03K | 30889634 | MI_only | trans | 57 | Midwest |
| Chr05K | 45117640 | TX_only | trans | 51 | Midwest |
| Chr05K | 64462169 | TX_only | trans | 68 | Midwest |
| Chr07K | 50270854 | TX_only | trans | 52 | Midwest |
| Chr07K | 51042263 | TX_only | trans | 82 | Midwest |
| Chr09N | 78131723 | MI_only | trans | 118 | Midwest |
| Chr01N | 41198087 | MI_only | trans | 57 | Atlantic |
| Chr03K | 3439055 | MI_only | trans | 82 | Atlantic |
| Chr03N | 7790323 | TX_only | trans | 233 | Atlantic |
| Chr04K | 33016371 | MI_only | trans | 52 | Atlantic |
| Chr05K | 40601001 | MI_only | trans | 66 | Atlantic |
| Chr05K | 47444048 | MI_only | trans | 67 | Atlantic |
| Chr06N | 19809737 | MI_only | trans | 57 | Atlantic |
| Chr07N | 23489614 | TX_only | trans | 56 | Atlantic |
| Chr08N | 45567500 | MI_only | trans | 752 | Atlantic |
| Chr09K | 67315327 | TX_only | trans | 89 | Atlantic |

Supplementary Table 7. Overlap between cis-regulated genes in the allele-specific expression analysis and the eQTL maps in the Gulf, Atlantic, and Midwest populations, in the Michigan and Texas gardens.

| Habitat | Population | # overlapping genes |
|---|---|---|

| | | |
|---|---|---|
| Michigan_$F_1$ | Midwest | 2261 |
| Michigan _$F_1$ | Gulf | 4734 |
| Michigan _$F_1$ | Atlantic | 3797 |
| Texas_$F_1$ | Midwest | 1585 |
| Texas _$F_1$ | Gulf | 5566 |
| Texas _$F_1$ | Atlantic | 3658 |

Supplementary Table 8. Minor allele frequencies for static and dynamic cis and trans eQTLs in all populations and habitats.

| pop | presence | type | MAF |
|---|---|---|---|
| Atlantic | GxE | cis | 0.24 |
| Atlantic | GxE | trans | 0.109 |
| Atlantic | MI_only | cis | 0.235 |
| Atlantic | MI_only | trans | 0.113 |
| Atlantic | TX_and_MI | cis | 0.254 |
| Atlantic | TX_and_MI | trans | 0.113 |
| Atlantic | TX_only | cis | 0.229 |
| Atlantic | TX_only | trans | 0.103 |
| Gulf | GxE | cis | 0.219 |
| Gulf | GxE | trans | 0.102 |
| Gulf | MI_only | cis | 0.207 |
| Gulf | MI_only | trans | 0.102 |
| Gulf | TX_and_MI | cis | 0.248 |
| Gulf | TX_and_MI | trans | 0.0945 |
| Gulf | TX_only | cis | 0.21 |
| Gulf | TX_only | trans | 0.104 |
| Midwest | GxE | cis | 0.28 |
| Midwest | GxE | trans | 0.0959 |
| Midwest | MI_only | cis | 0.283 |
| Midwest | MI_only | trans | 0.1 |
| Midwest | TX_and_MI | cis | 0.297 |
| Midwest | TX_and_MI | trans | 0.118 |
| Midwest | TX_only | cis | 0.254 |
| Midwest | TX_only | trans | 0.0883 |

Supplementary Table 9. Candidate genes from the leaf phosphorus concentration GWAS in each site and population, with their annotations and whether they overlap with mapped cis and trans eQTL, attached as Bergetal_TableS9.csv.

Supplementary Table 10. Leaf phosphorus candidate genes regulated by the Gulf trans eQTL, which acts as a regulatory hotspot in the Texas garden, attached as Bergetal_TableS10.csv.

Supplementary Figures

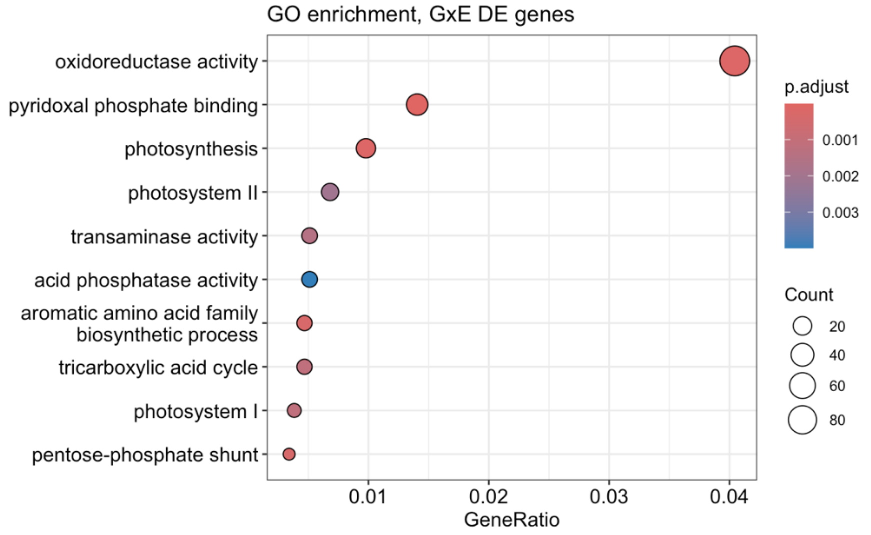

Figure S1. **GxE DE Gene GO Terms**. GO enrichment terms in the significant differentially expressed genes with genotype x environment interactions. Model = expression ~ genotype + environment + genotype:environment.

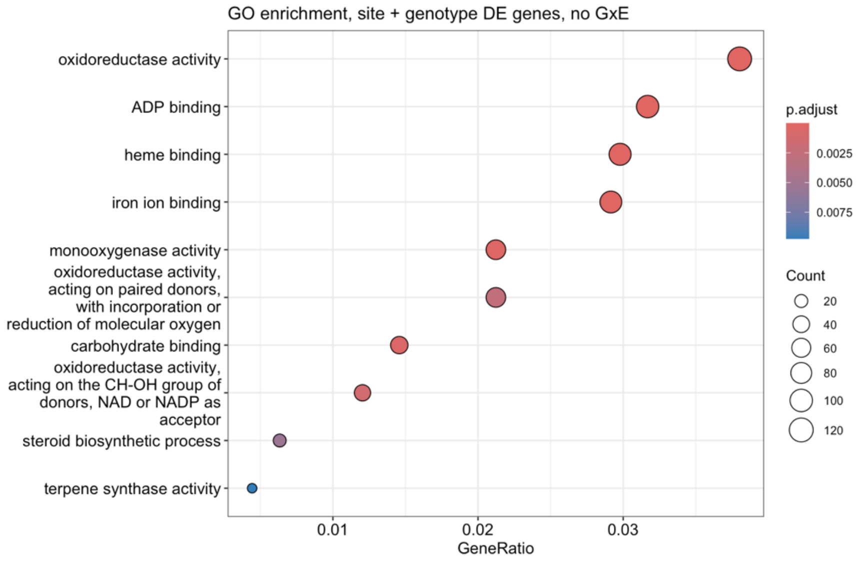

Figure S2. **Site and genotype DE gene GO terms**. GO enrichment terms in the significant differentially expressed genes whose expression was significantly shaped by site and genotype, but not their interaction. Model = expression ~ genotype + environment + genotype:environment.

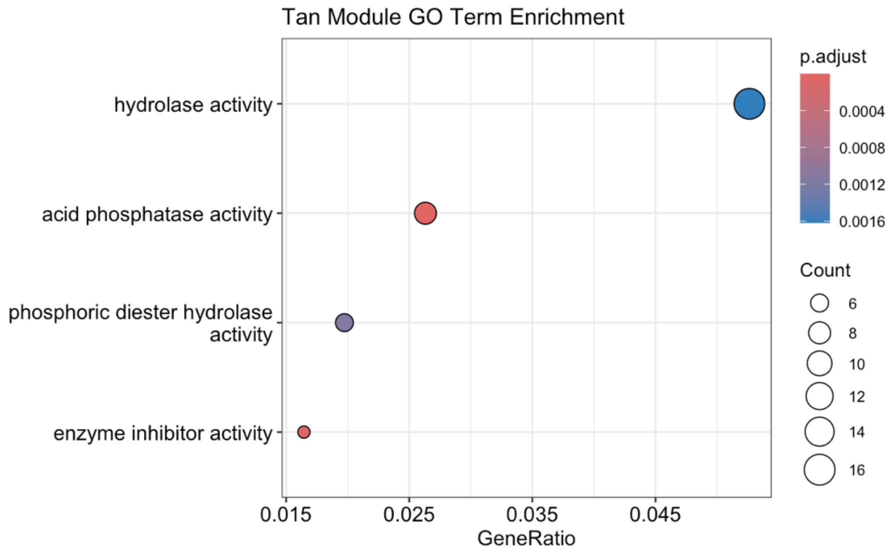

Figure S3. **Tan Module GO Terms**. GO enrichment terms from the tan module in weighted gene coexpression network analysis of gene expression in AP13 and DAC6 in Michigan and Texas common gardens.

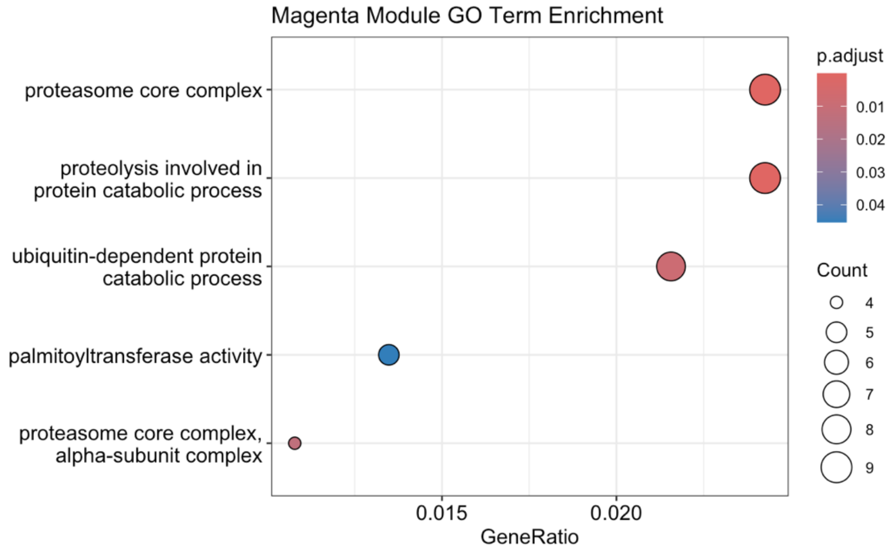

Figure S4. **Magenta Module GO Terms**. GO enrichment terms from the magenta module in weighted gene coexpression network analysis of gene expression in AP13 and DAC6 in Michigan and Texas common gardens.

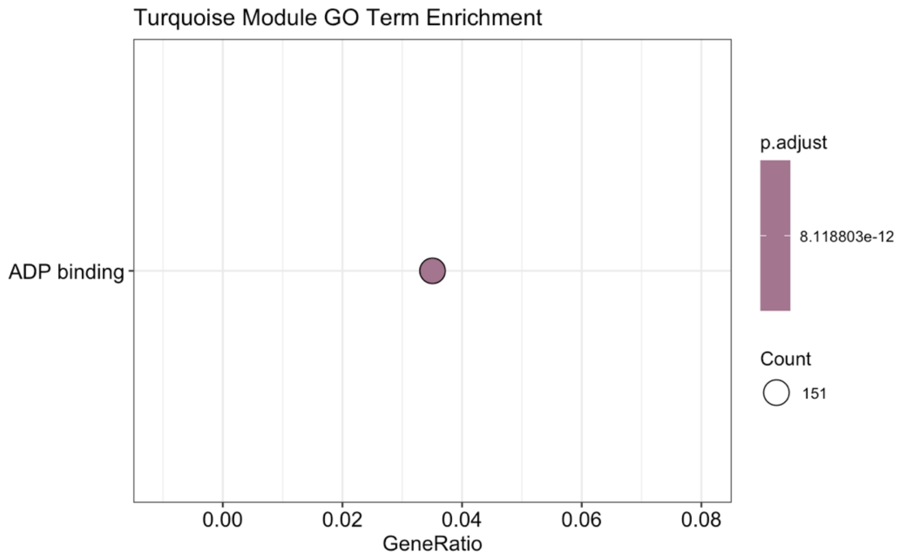

Figure S5. **Turquoise Module GO Term**. GO enrichment term from the turquoise module in weighted gene coexpression network analysis of gene expression in AP13 and DAC6 in Michigan and Texas common gardens.

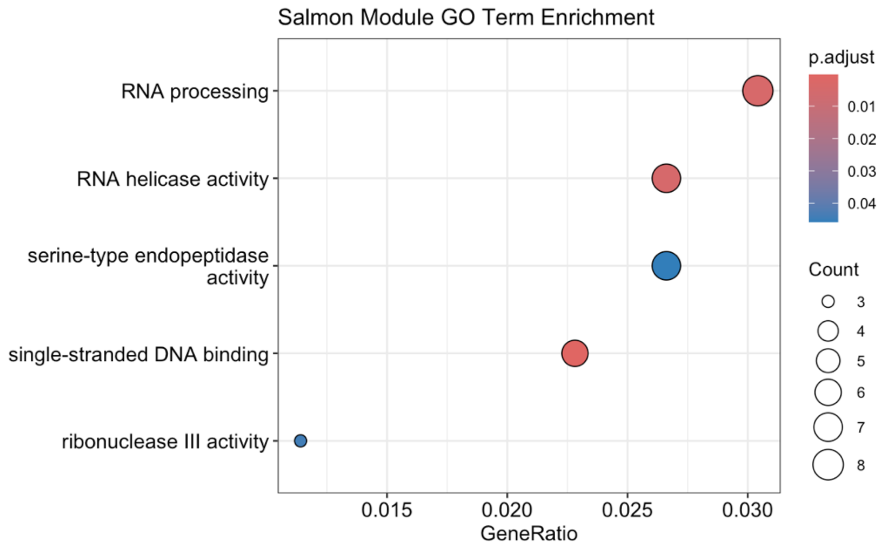

Figure S6. **Salmon Module GO Terms**. GO enrichment terms from the salmon module in weighted gene coexpression network analysis of gene expression in AP13 and DAC6 in Michigan and Texas common gardens.

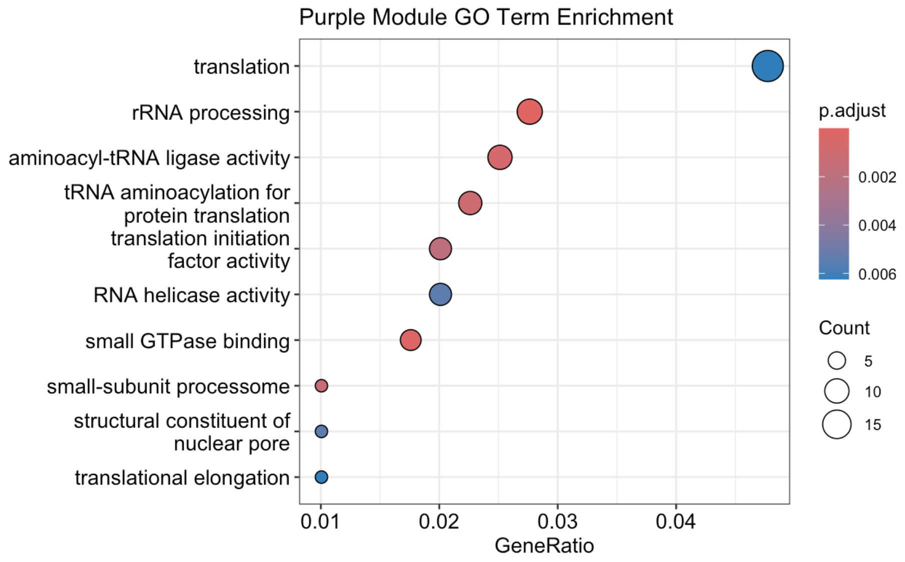

Figure S7. **Purple Module GO Terms**. GO enrichment terms from the purple module in weighted gene coexpression network analysis of gene expression in AP13 and DAC6 in Michigan and Texas common gardens.

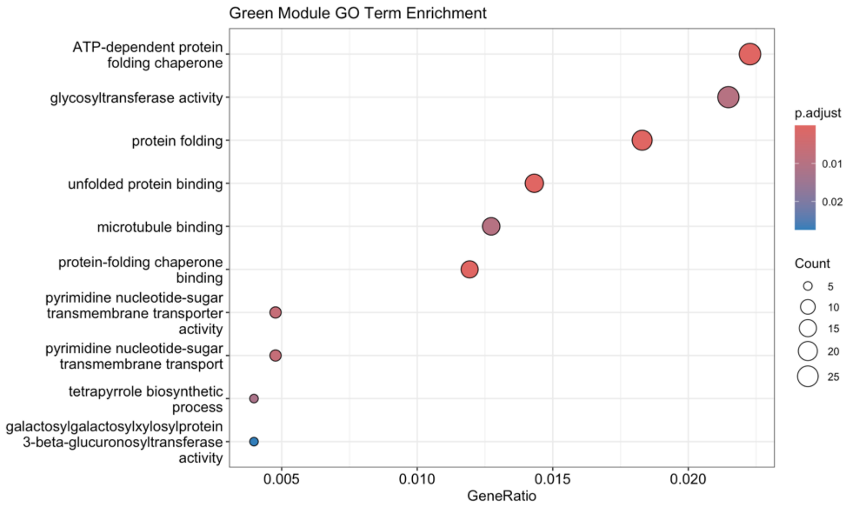

Figure S8. **Green Module GO Terms.** GO enrichment terms from the green module in weighted gene coexpression network analysis of gene expression in AP13 and DAC6 in Michigan and Texas common gardens.

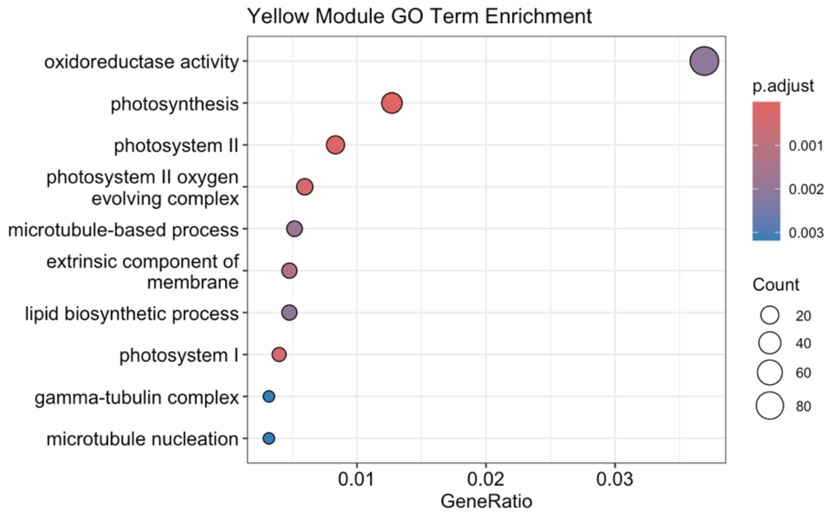

Figure S9. **Yellow Module GO Terms.** GO enrichment terms from the yellow module in weighted gene coexpression network analysis of gene expression in AP13 and DAC6 in Michigan and Texas common gardens.

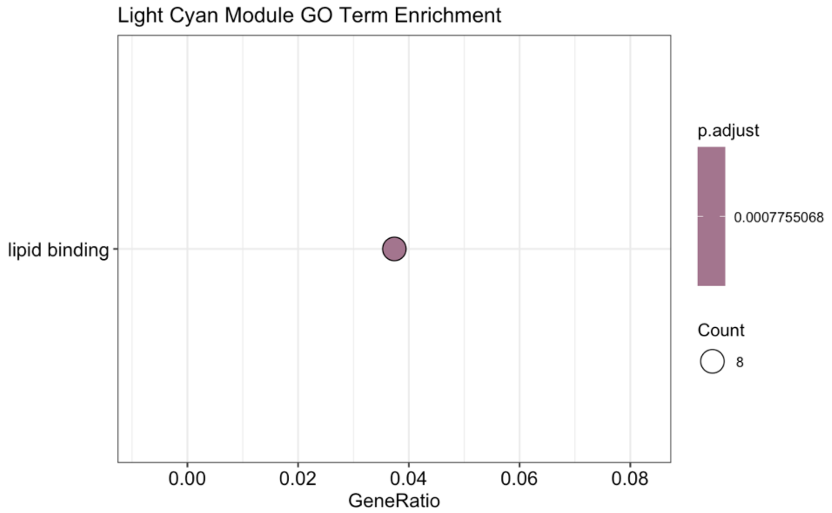

Figure S10. **Light Cyan Module GO Term.** GO enrichment term from the yellow module in weighted gene coexpression network analysis of gene expression in AP13 and DAC6 in Michigan and Texas common gardens.

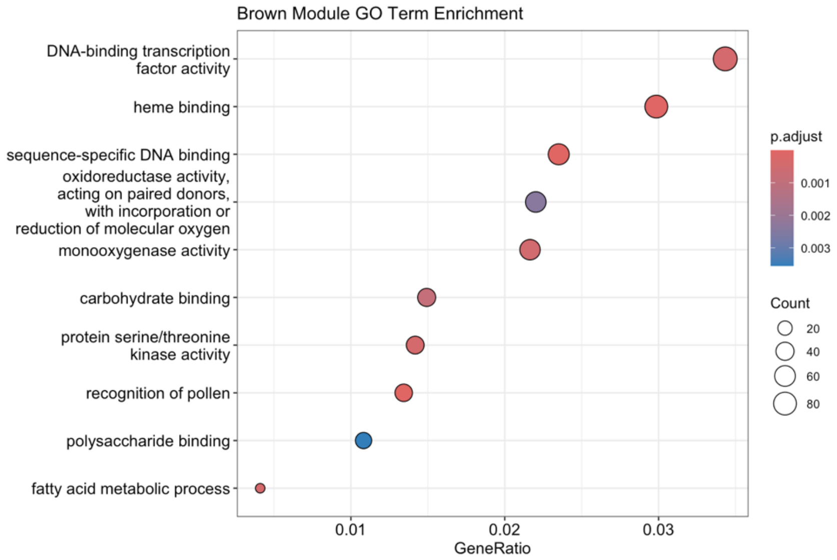

Figure S11. **Brown Module GO Terms.** GO enrichment terms from the brown module in weighted gene coexpression network analysis of gene expression in AP13 and DAC6 in Michigan and Texas common gardens.

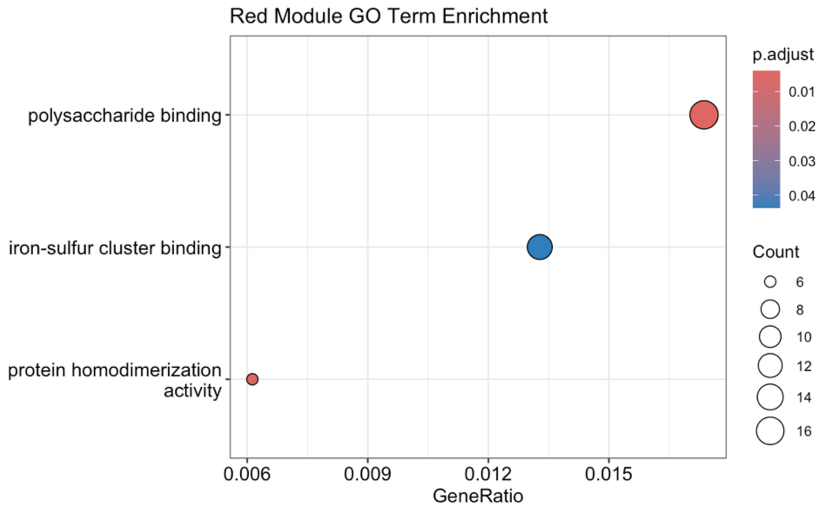

Figure S12. **Red Module GO Terms.** GO enrichment terms from the red module in weighted gene coexpression network analysis of gene expression in AP13 and DAC6 in Michigan and Texas common gardens.

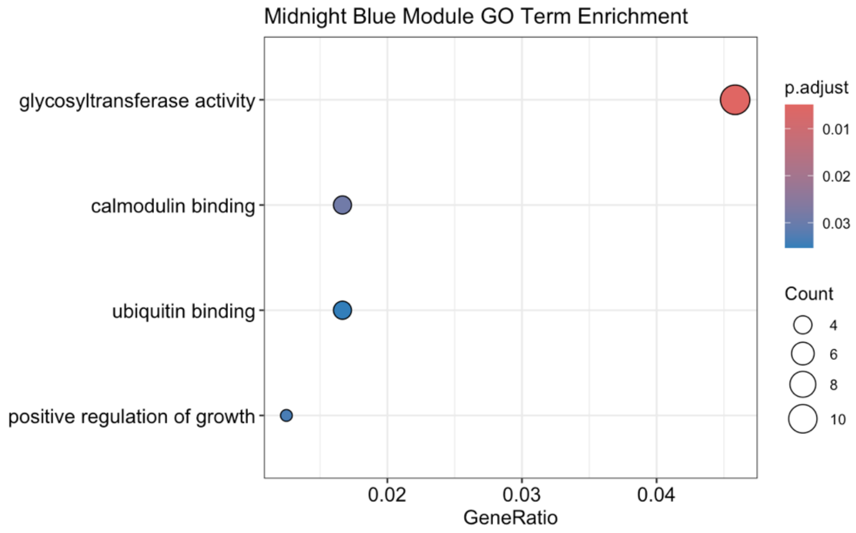

Figure S13. **Midnight Module GO Terms.** GO enrichment terms from the midnight blue module in weighted gene coexpression network analysis of gene expression in AP13 and DAC6 in Michigan and Texas common gardens.

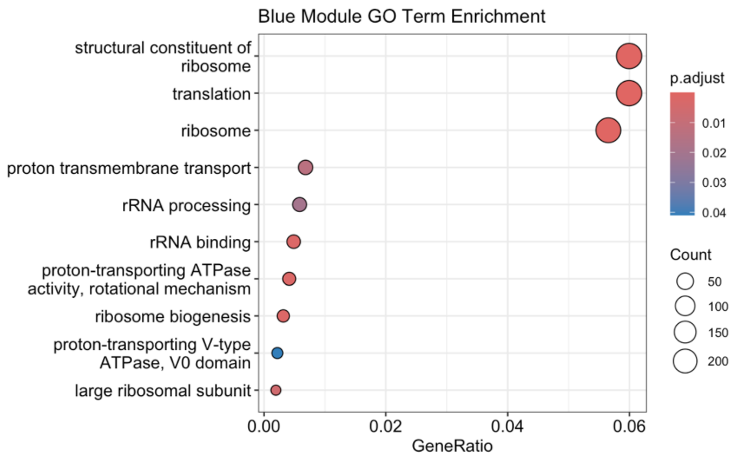

Figure S14. **Blue Module GO Terms.** GO enrichment terms from the blue module in weighted gene coexpression network analysis of gene expression in AP13 and DAC6 in Michigan and Texas common gardens.

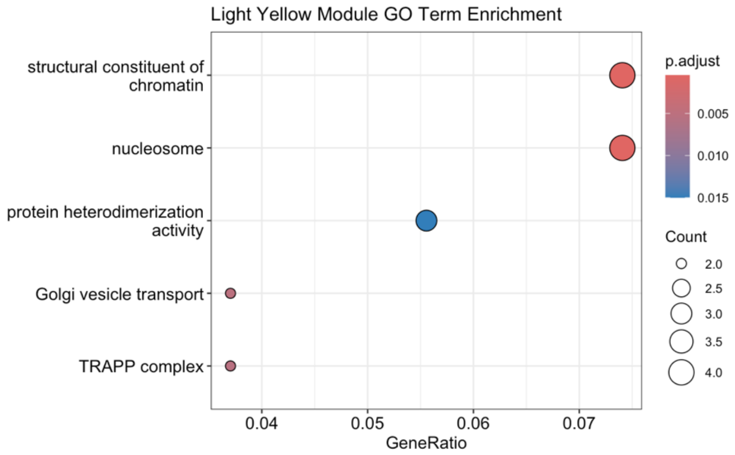

Figure S15. **Light yellow Module GO Terms.** GO enrichment terms from the blue module in weighted gene coexpression network analysis of gene expression in AP13 and DAC6 in Michigan and Texas common gardens.

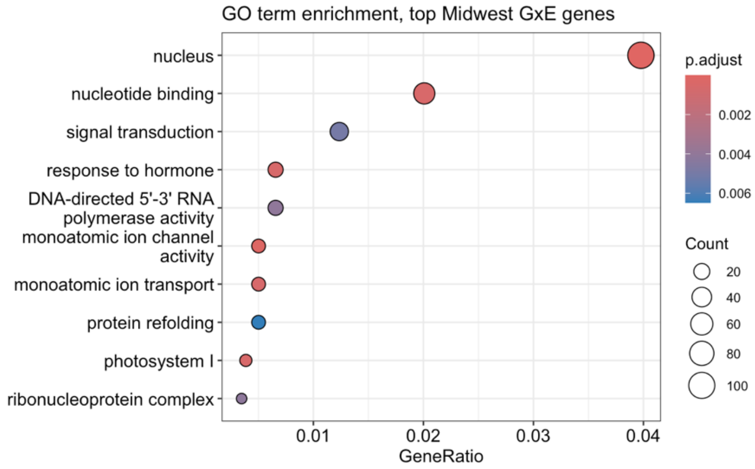

Figure S16. Midwest population top **GxE Gene GO Terms**. GO enrichment terms for the top 10% of genes with a change in expression between MI and TX sites in the Midwest population of the diversity panel.

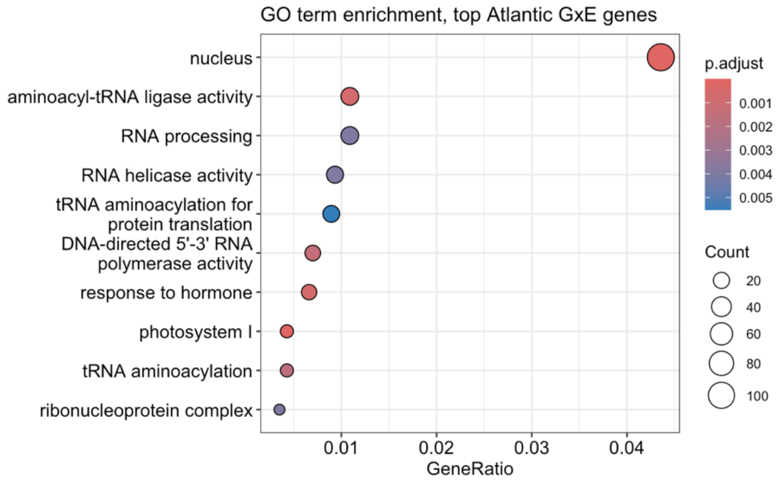

Figure S17. Atlantic population top **GxE Gene GO Terms**. GO enrichment terms for the top 10% of genes with a change in expression between MI and TX sites in the Atlantic population of the diversity panel.

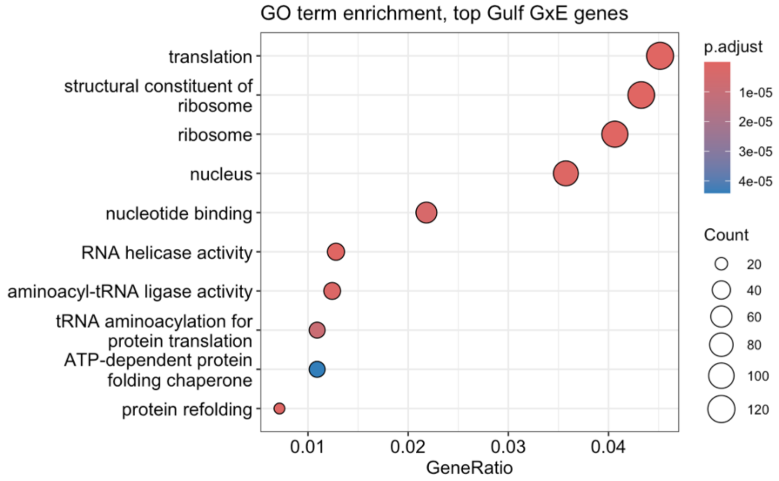

Figure S18. Gulf population top **GxE Gene GO Terms**. GO enrichment terms for the top 10% of genes with a change in expression between MI and TX sites in the Gulf population of the diversity panel.

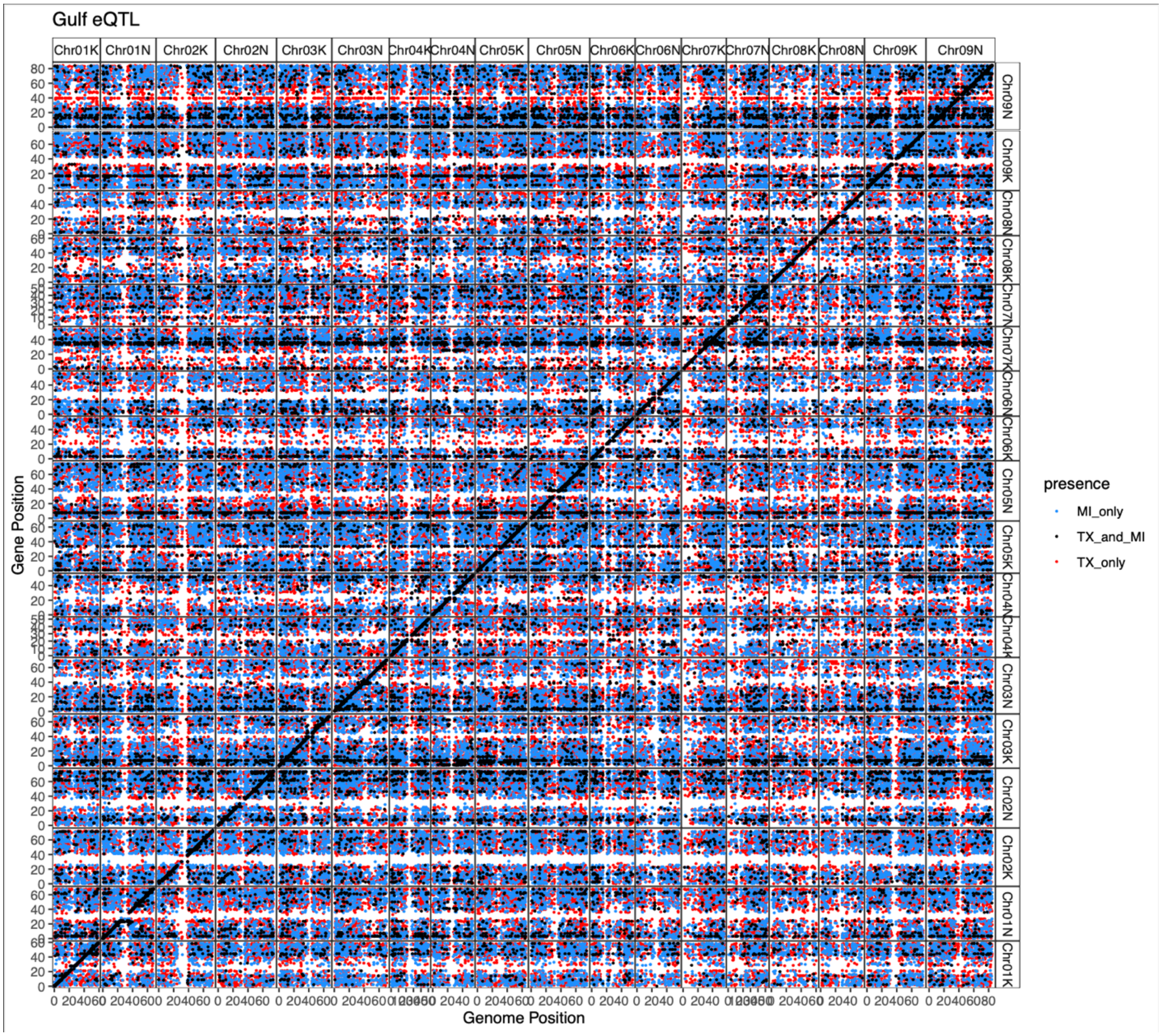

Figure S19. **Static and dynamic cis and trans eQTL in the Gulf population.** Points on the diagonal line are cis-eQTLs; off-diagonal points are trans-eQTLs. Dynamic eQTLs, which are specific to habitat, are in blue and red. Blue points were mapped only in Michigan, red points are mapped only in Texas. Black points are static eQTLs, present in both MI and TX

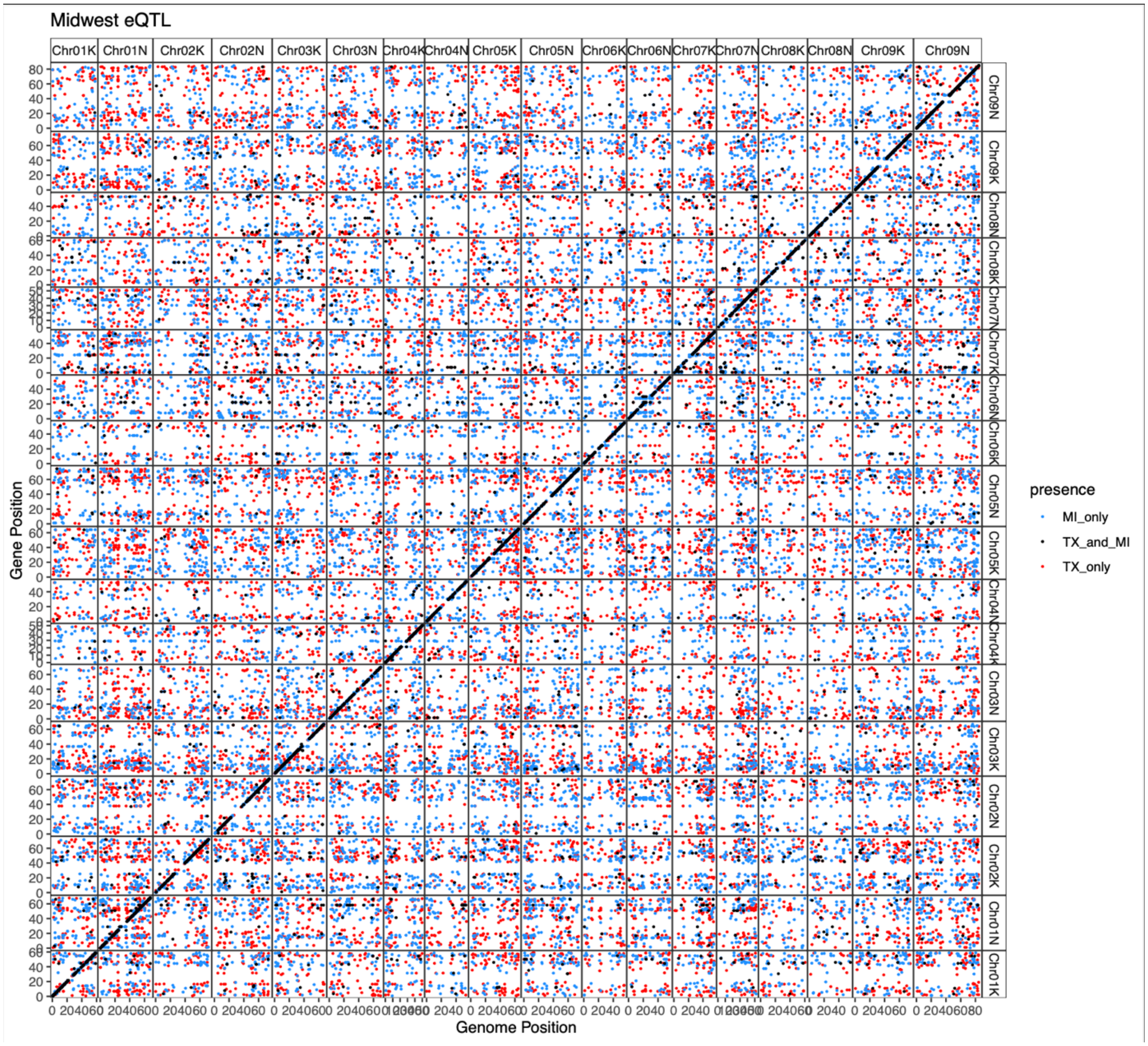

Figure S20. **Static and dynamic cis and trans eQTL in the Midwest population.** Points on the diagonal line are cis-eQTLs; off-diagonal points are trans-eQTLs. Dynamic eQTLs, which are specific to habitat, are in blue and red. Blue points were mapped only in Michigan, red points are mapped only in Texas. Black points are static eQTLs, present in both MI and TX

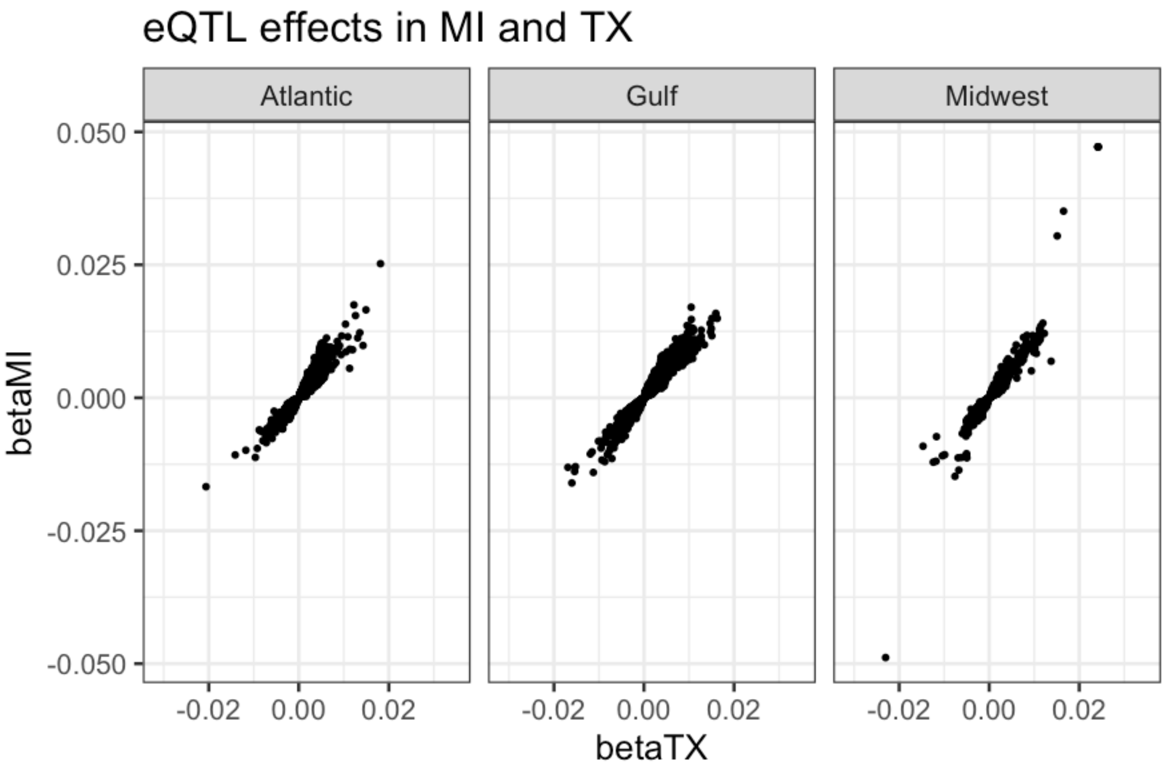

**Figure S21.** Effects (beta) of eQTL in Texas and Michigan, in all three populations of the diversity panel.

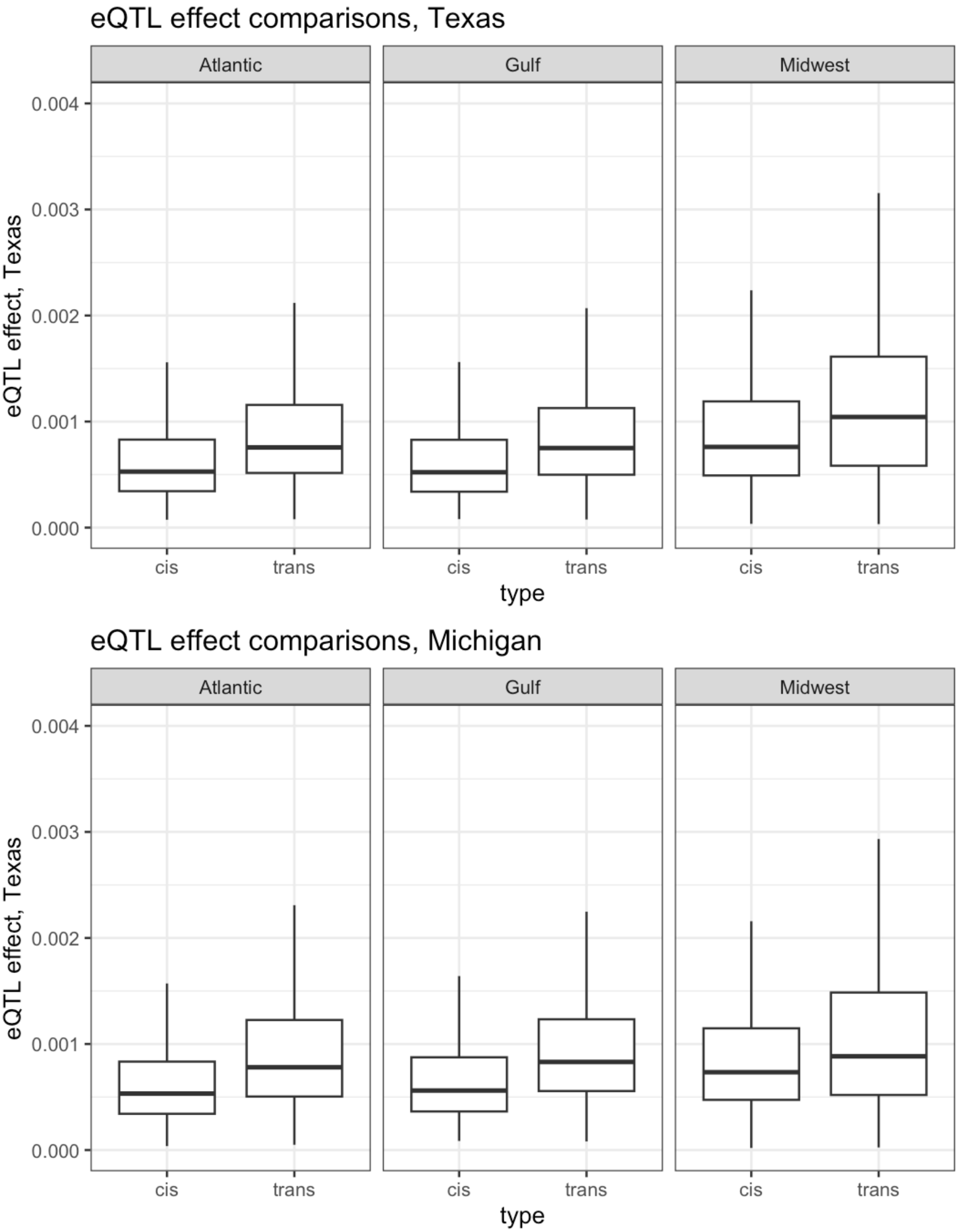

Figure S22. eQTL effects in MI and TX in all three populations of the diversity panel. p-value < 2.2e-16 in all t.test comparisons between betas of cis and trans eQTL.

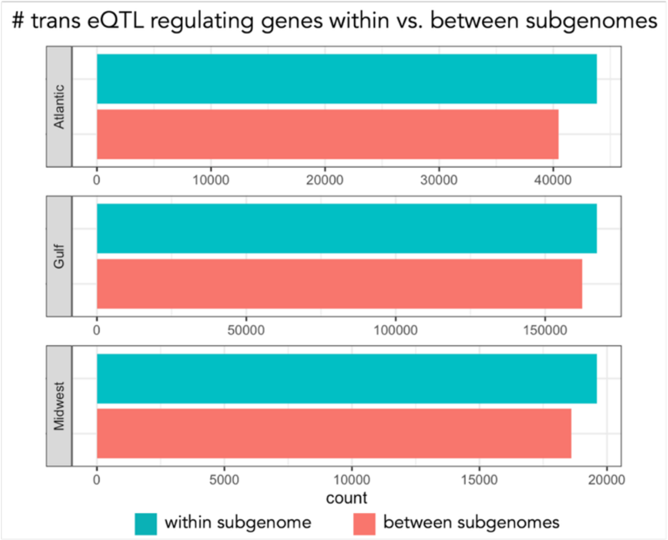

**Figure S23.** Comparison of number of trans-eQTL regulating expression of genes within vs. between subgenomes

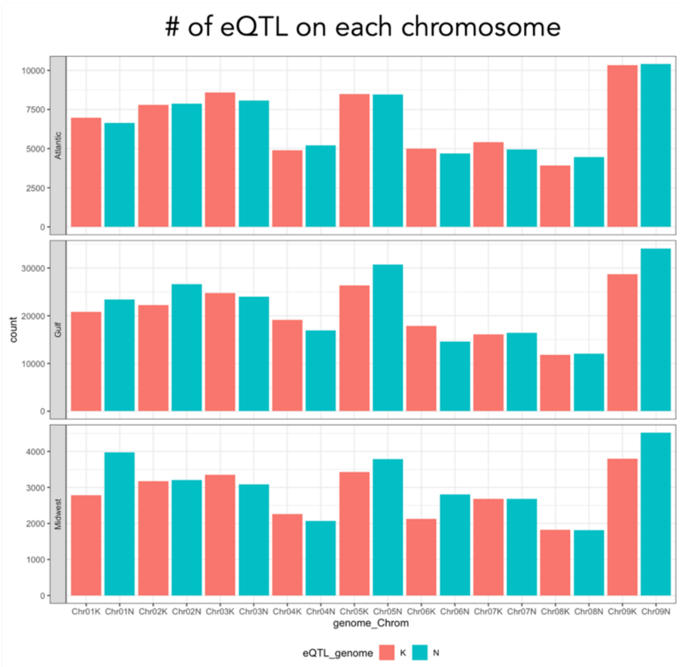

**Figure S24.** Counts of eQTL on each chromosome in the Midwest, Gulf, and Atlantic populations

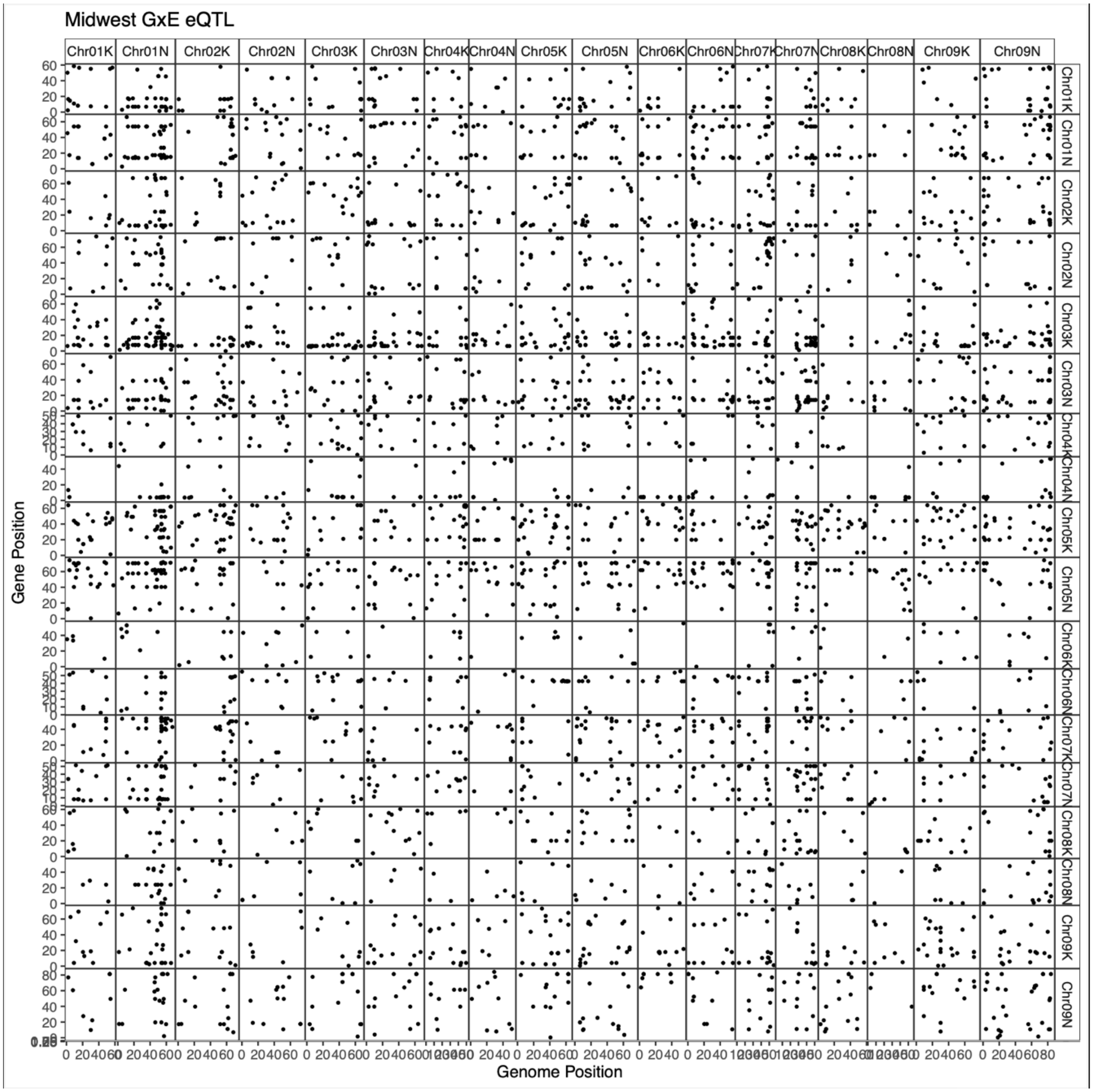

Figure S25. **Genotype x Environment eQTLs in the Midwest population.** Each point represents an eQTL that significantly predicted a difference in gene expression between the Michigan and Texas common gardens. Points on the diagonal line are cis-eQTLs; off-diagonal points are trans-eQTLs.

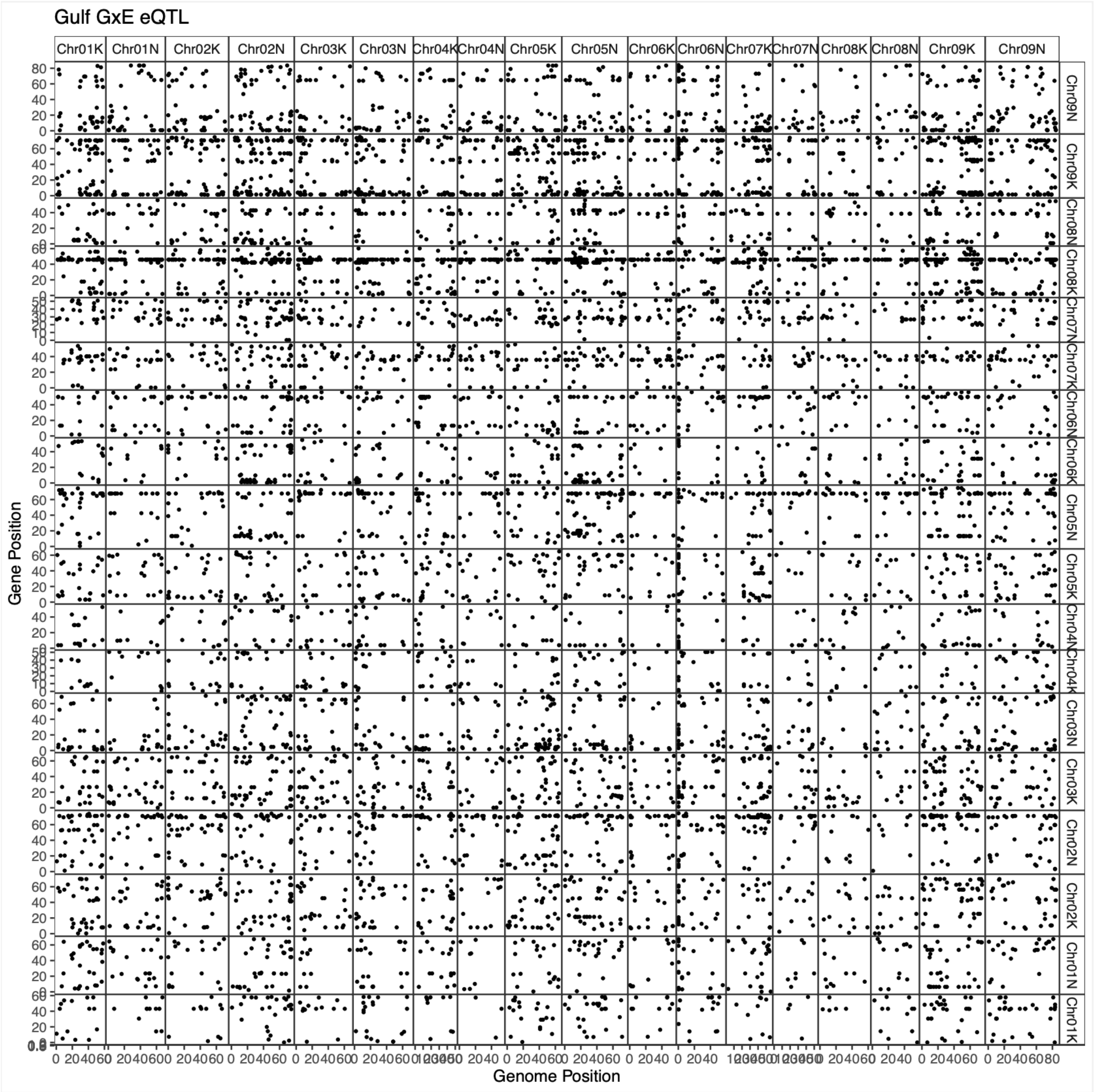

Figure S26. **Genotype x Environment eQTLs in the Gulf population.** Each point represents an eQTL that significantly predicted a difference in gene expression between the Michigan and Texas common gardens. Points on the diagonal line are cis-eQTLs; off-diagonal points are trans-eQTLs.

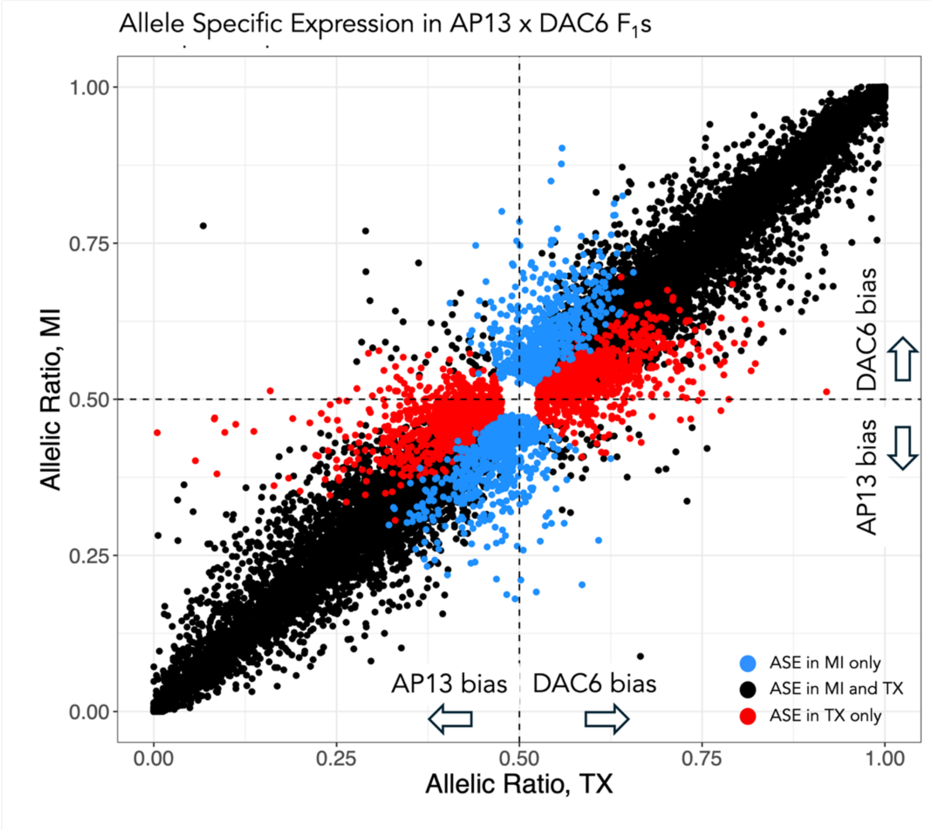

Figure S27. **Allele-specific expression in AP13xDAC6 $F_1$'s suggests *cis*-regulation of gene expression is stable across sites.** Ratio of AP13 and DAC6 alleles in genes with significant allele-specific expression in the Michigan and TX sites. Color indicates whether the bias was site-specific (MI only or TX only) or mapped in both sites.

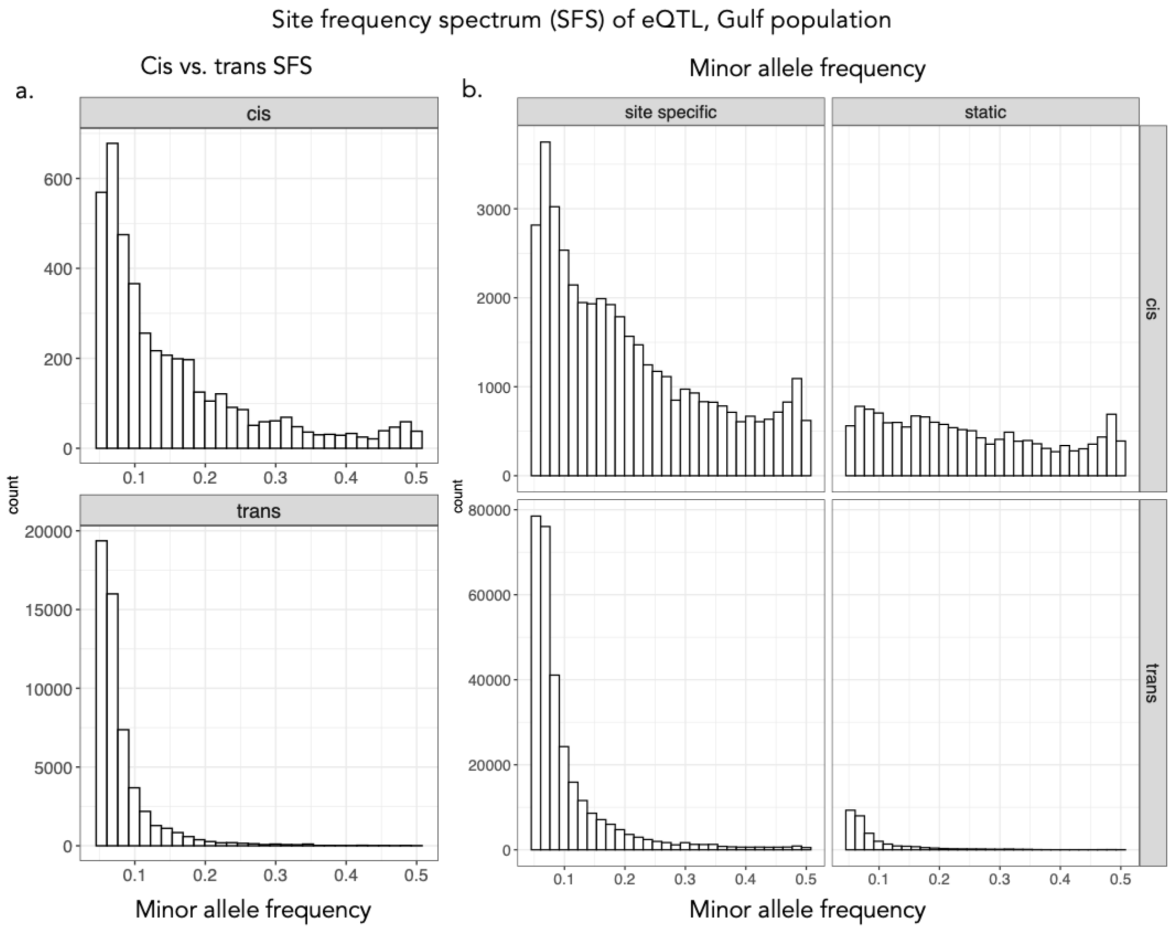

Figure S28. **Site frequency spectrum comparisons of Gulf eQTL.** a) Site frequency spectrum of cis and trans eQTL in the Gulf population of the diversity panel, with eQTL effects in the top quartile. b) Site frequency spectrum of cis and trans eQTL that are site-specific (mapped in only TX or MI) and static (mapped in both TX and MI) in the Atlantic population of the switchgrass diversity panel.

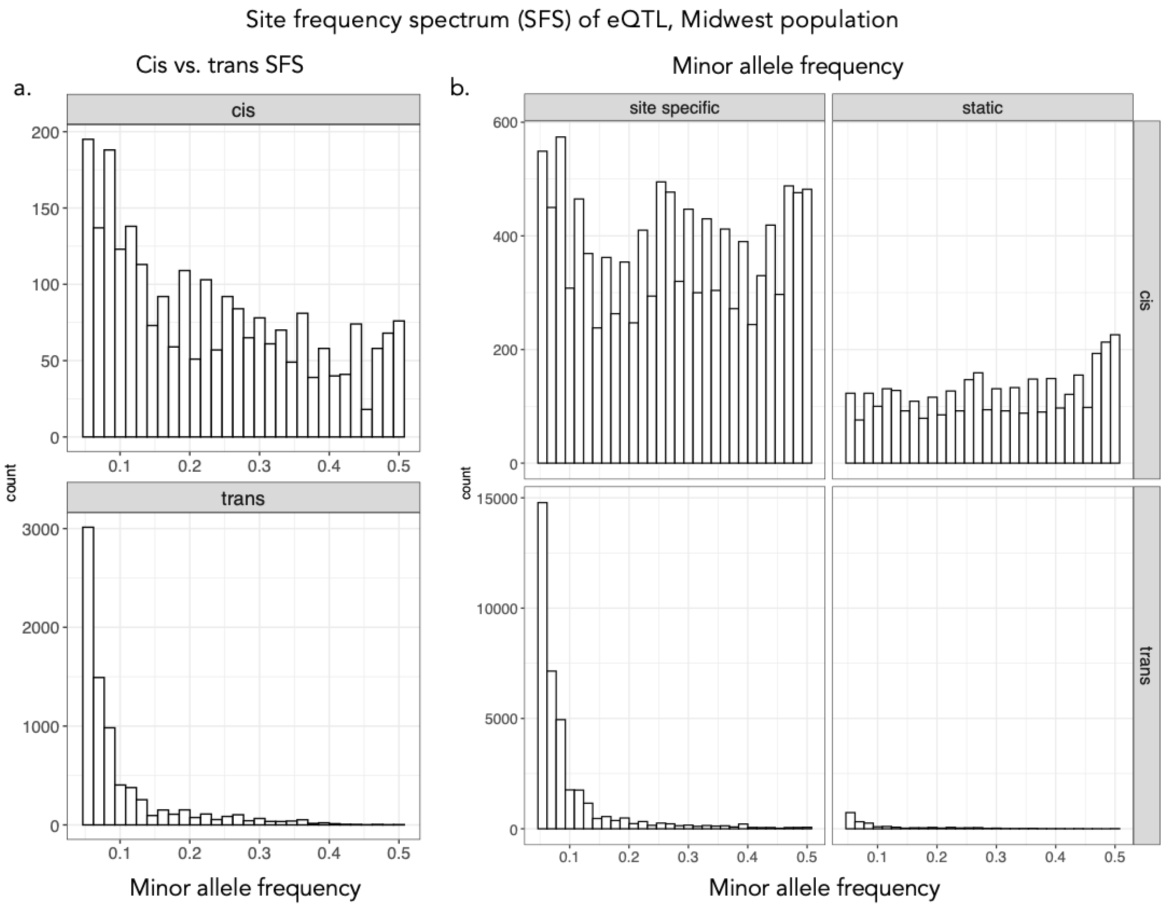

Figure S29. **Site frequency spectrum comparisons of Midwest eQTL.** a) Site frequency spectrum of cis and trans eQTL in the Midwest population of the diversity panel, with eQTL effects in the top quartile. b) Site frequency spectrum of cis and trans eQTL that are site-specific (mapped in only TX or MI) and static (mapped in both TX and MI) in the Atlantic population of the switchgrass diversity panel.

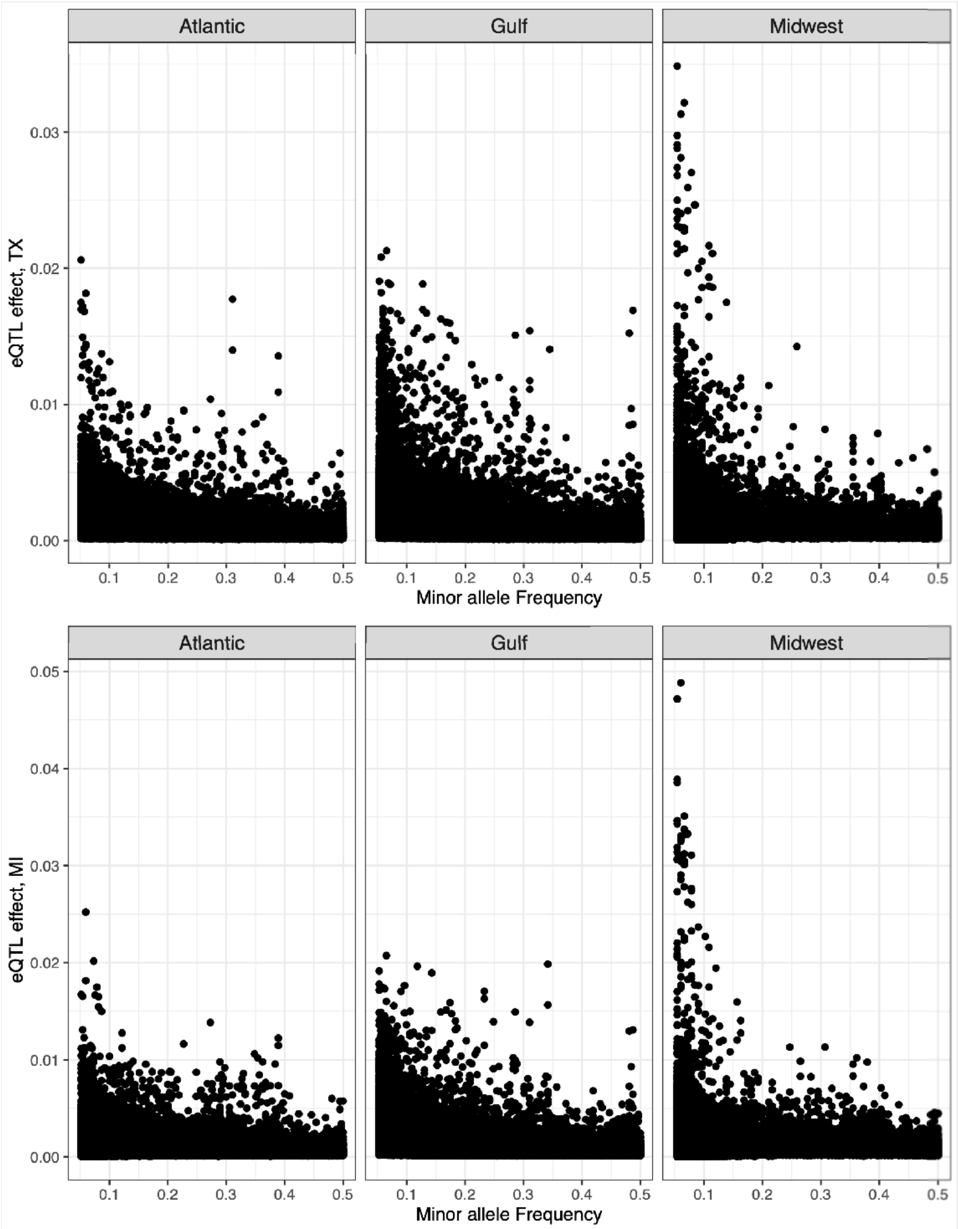

Fig. S30. **eQTL with lower minor allele frequencies have higher absolute effect sizes.** Relationship between minor allele frequency and eQTL effect size in Texas (top) and Michigan (bottom), in all three populations of the swithgrass diversity panel.

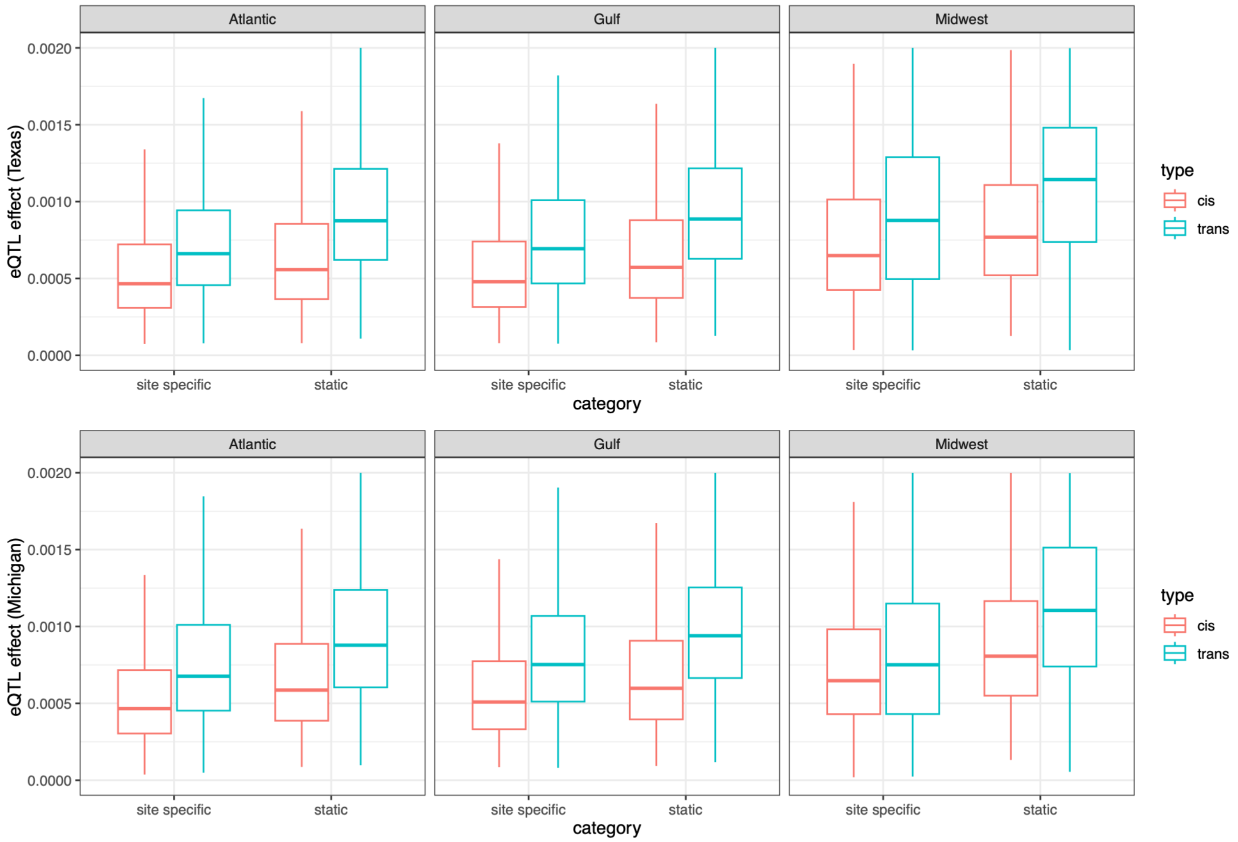

Fig. S31. Comparison of eQTL effect sizes among site-specific and static cis and trans eQTL in MI and TX, split by population

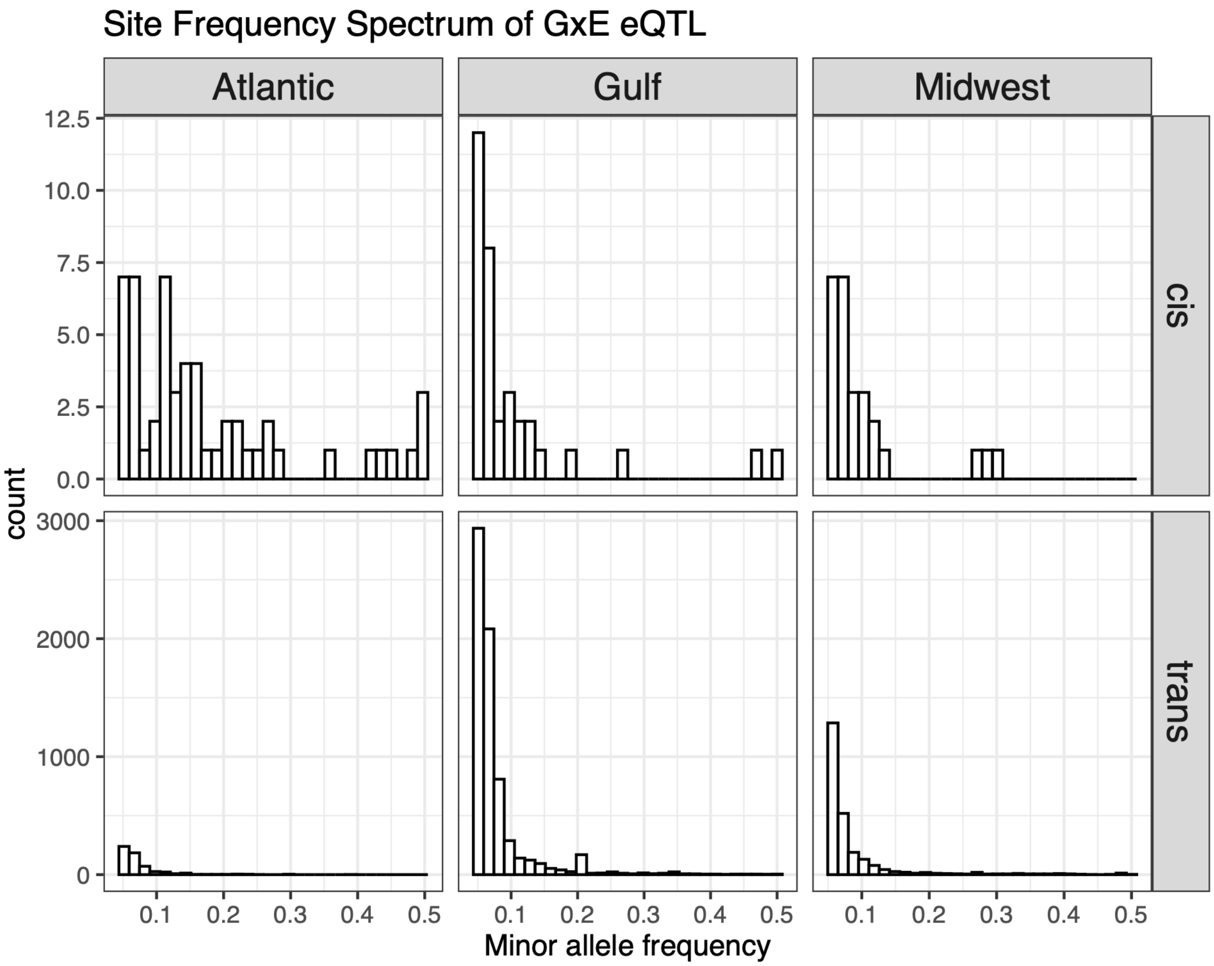

Figure S32. **Site frequency spectrum comparisons of GxE eQTL.** a) Site frequency spectrum of cis and trans eQTL that predicted a change in gene expression between the MI and TX sites (GxE eQTL), for all three populations in the switchgrass diversity panel.

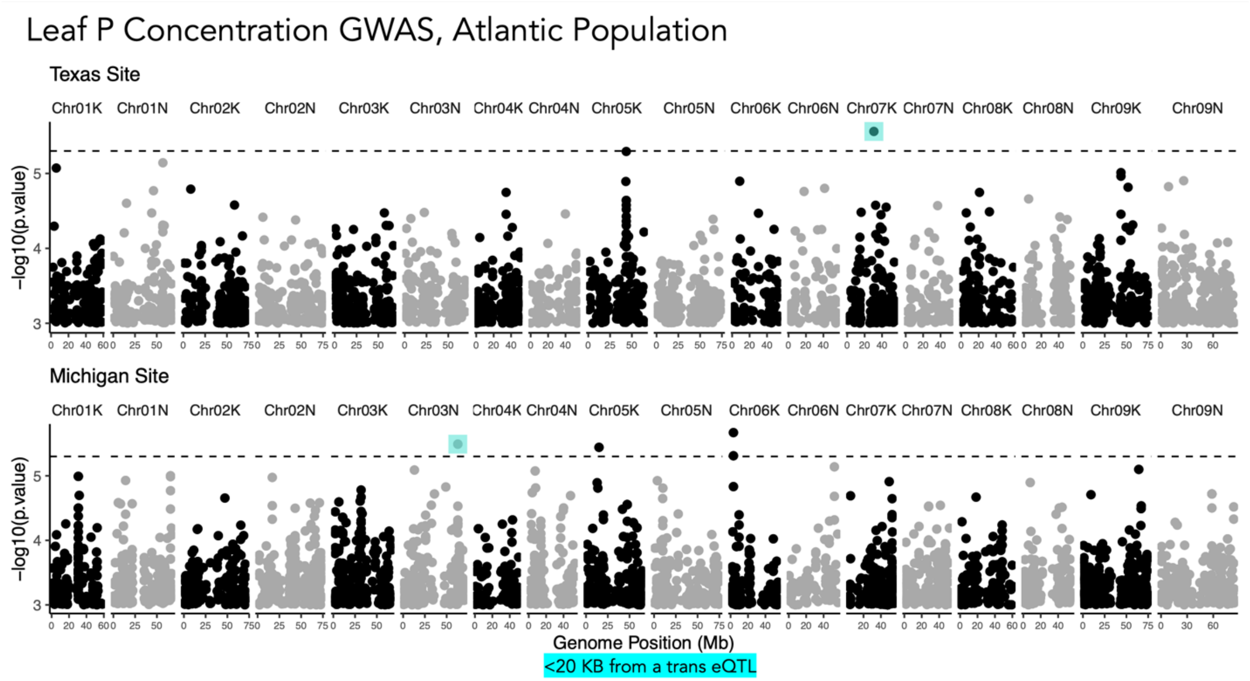

Figure S33. Switchgrass Leaf Phosphorus concentration GWAS, Atlantic population at the Texas and Michigan sites. Highlighted points are candidate SNPs within <20Kb of a trans eQTL from Figure 3.

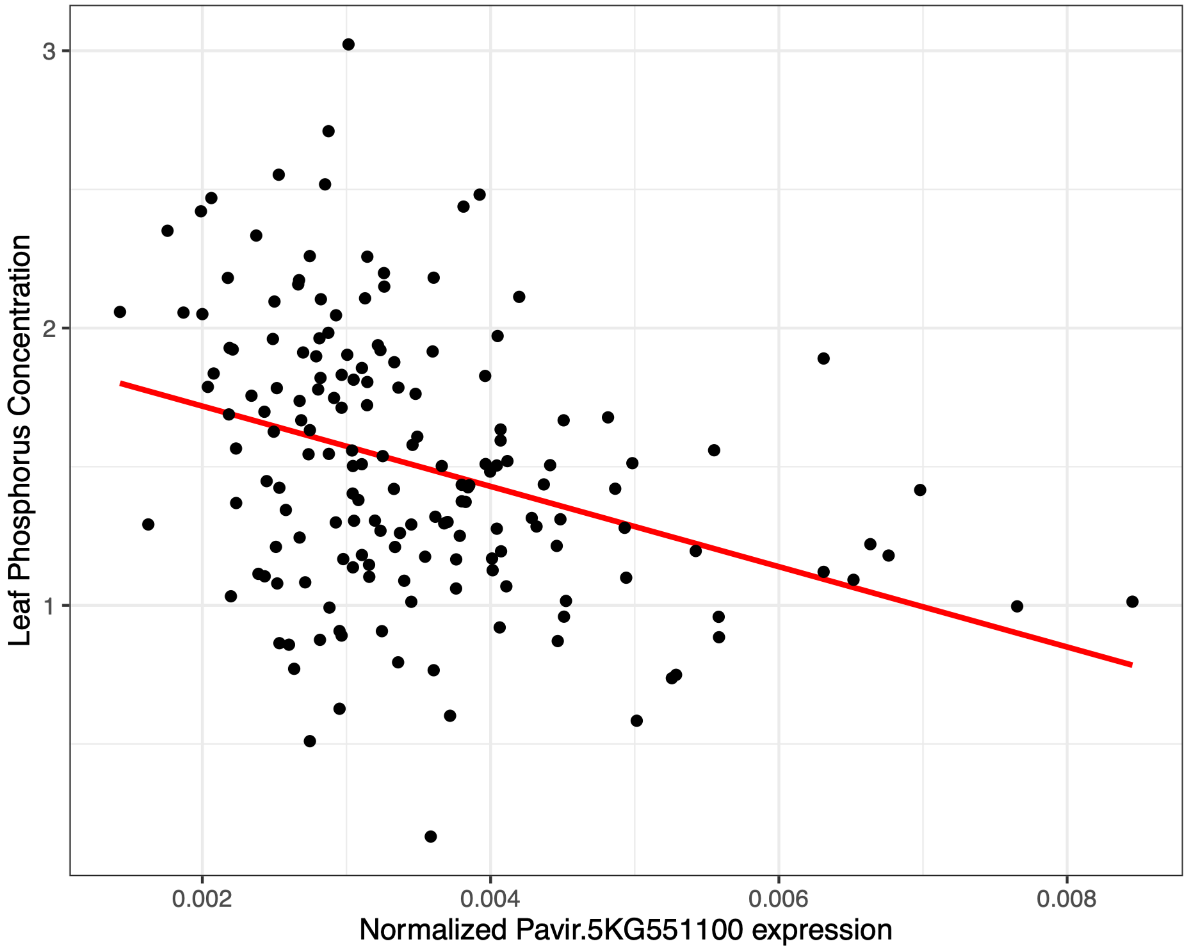

Figure S34. Normalized expression of Pavir.5KG551100 significantly predicts leaf phosphorus concentration in the Gulf population at the Texas site (estimate = -165.6, Adjusted p-value = 0.018, Adjusted $R^2$ = 0.141).

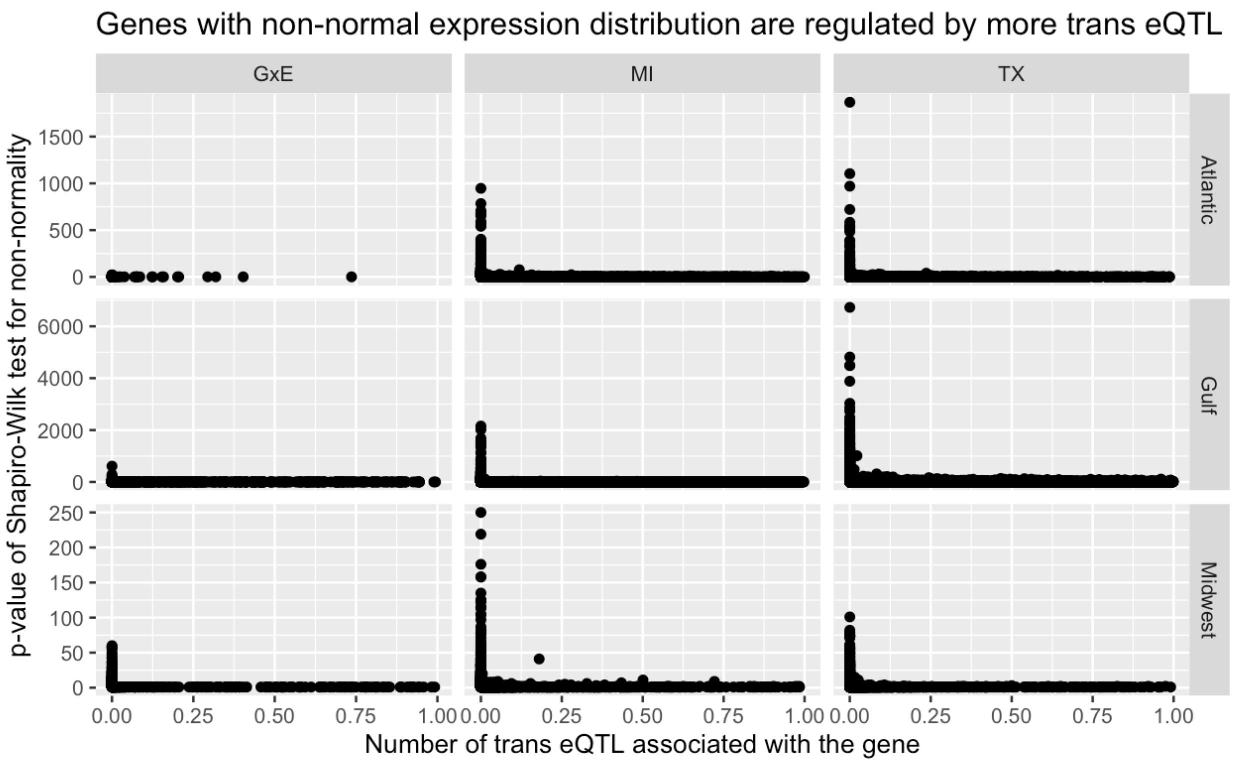

**Figure S35. Genes with non-normal expression distribution are regulated by more trans eQTL.**
